# Coreset-based logistic regression for atlas-scale single-cell and spatial omics analyses

**DOI:** 10.1101/2025.06.04.655742

**Authors:** Danrong Li, Van Hoan Do, Young Kun Ko, Stefan Canzar

**Affiliations:** Department of Computer Science and Engineering, The Pennsylvania State University, University Park, PA 16802, United States; Center for Applied Mathematics and Informatics, Le Quy Don Technical University, Hanoi, Vietnam; Faculty of Informatics and Data Science, University of Regensburg, Regensburg, Germany

## Abstract

Logistic regression provides a versatile and interpretable framework for analyzing single-cell and spatial omics data. It has been shown to outperform more complex models on key association tasks, such as marker gene identification, and on predictive tasks, such as cell type annotation. Logistic regression has also been used effectively for quantifying spatial proximity between cell types and has demonstrated competitive performance in predicting tissue phenotypes. As increasingly large single-cell atlases become publicly available, training logistic regression models on these datasets offers opportunities for building accurate and robust reference models but also poses major computational challenges. Here, we adapt and extend the theory of coresets for logistic regression. Specifically, we compute a previously introduced classification complexity measure using linear programming to identify omics datasets that admit coresets, i.e. random subsets of cells that preserve logistic loss. Our main theoretical finding is that this complexity measure is close to one for PCA-transformed single-cell datasets. Moreover, we prove that logistic loss is preserved under this linear transformation, suggesting a universal sampling scheme that requires only a constant number of cells per cell type to obtain accurate representations of the data. Experiments across several cell atlases demonstrate that these theoretical guarantees translate to accurate and scalable transfer of cell types and spatial niches, outperforming more complex models, including pre-trained language models and graph isomorphism networks.

## 1 Introduction

Logistic regression offers a simple yet powerful and interpretable framework for the analysis of single-cell and spatial omics data. Despite its conceptual simplicity, it has been shown to outperform more complex machine learning models in key association tasks such as marker gene identification [1], as well as in predictive applications including cell type annotation [2]. Moreover, logistic regression has been successfully applied to quantify spatial relationships between cell types [3] and to predict tissue-level phenotypes with competitive accuracy, even when compared to graph-based deep learning models such as graph convolutional networks (GCNs) and graph isomorphism networks (GINs) that explicitly operate on spatial cell-cell graphs, especially in straightforward classification settings such as predicting tumor grade in breast cancer [4].

Among these applications, cell type annotation remains a central task in single-cell omics analysis. Machine learning and deep learning models trained on publicly available single-cell datasets are increasingly being used for this purpose. In particular, foundation models such as scFoundation [5] and scGPT [6] which are pre-trained on vast amounts of unlabeled single-cell RNA-seq data and then fine-tuned on a smaller, task-specific dataset have demonstrated enormous success in the predictive analysis of single-cell data. However, experiments have shown that classic machine learning methods such as logistic regression have an advantage over large pre-trained models on relatively simple tasks, including cell type annotation [2]. In fact CellTypist [7], an immune cell atlas of myeloid and lymphoid lineages in human tissue was developed based on a logistic regression model trained on more than 700,000 cells from 19 different studies covering 20 different tissue types. For convergence and storage reasons, the training data was divided into many small batches on which training by stochastic gradient descent was carried out one at a time. While this strategy achieved a substantial learning speed-up compared to conventional logistic regression, it provides no guarantees on the quality of the resulting solution.

In this work we propose an alternative approach based on the theory of coresets to train a logistic regression model on very large datasets efficiently and with theoretically guaranteed performance. In particular, we sketch a very large input matrix, that is, we randomly sample a small subset of cells (rows of the matrix), a so-called coreset [8, 9], so that the optimization of the much smaller problem instance yields a solution that is close to the optimal solution of the full problem. This method however does not work for all large datasets. Some datasets are known to be resilient to sampling [10]. This is captured by a previously introduced classification complexity measure *µ* which identifies single-cell datasets which admit coresets that preserve logistic loss and regression coefficients. In particular, if a dataset has a small *µ* value, it admits a small coreset depending on how small *µ* is [8, 10]. Note that this is a sufficient condition for coresets. There are datasets with bad *µ* value that empirically admit a good coreset, alas without theoretical guarantee.

To quantify dataset-specific sampling behavior, we implement a linear program (LP) [11] that computes the classification complexity for a range of single-cell datasets. Corroborating our empirical observations, our main theoretical result shows that this complexity measure approaches one for PCA-transformed single-cell data. Moreover, we prove that the logistic loss is preserved under this linear transformation, suggesting a universal sampling scheme in which only a constant number of cells per cell type are required to obtain accurate representations of the data.

In experiments on several cell atlases spanning different species and tissues, we compared our method to CellTypist and scMulan [12], a pre-trained language model, to demonstrate that our theoretical guarantees translate into accurate and scalable cell type label transfer across atlas-scale datasets. Furthermore, in spatial transcriptomics data, we showed that these guarantees similarly enable reliable transfer of spatial niches, i.e., localized communities of interacting cells, across tissue sections, outperforming more complex graph-based models such as graph isomorphism networks.

## 2 Methods

After formally introducing the problem setting, we first prove (Theorem 1) that the logistic loss is preserved under a rank-*k* approximation (e.g., PCA) and subsequent coreset construction, using a straightforward application of the triangle inequality. The coreset constructed in Theorem 1 depends on a complexity measure *µ*, which we show in our main theoretical result (Theorem 2) to be small for PCA-transformed single-cell data. Together, these findings suggest a universal sampling scheme in which only a constant number of cells per cell type are required to obtain accurate representations of the data (see Section 2.2).

### 2.1 Small complexity of PCA-transformed single-cell data

#### Problem Setting

Define a data matrix *H* ∈ ℝ^*n*×*d*^ and a label vector *y* ∈ {±1}^*n*^. The logistic regression problem seeks to minimize, over *β* ∈ ℝ^*d*^, the logistic loss

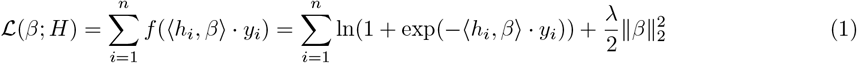

where ∀*i* ∈ [*n*], *h*_*i*_ represents the *i*-th row of *H* and *λ >* 0 is the *ℓ*_2_ regularization parameter. Define the optimal logistic regression coefficient as 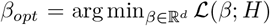. Finding such *β*_*opt*_ is costly, especially when both *n* and *d* can be really large. For biological data, in order to mitigate this issue, popular methods like CellTypist [7] use stochastic gradient descent (SGD) with randomly selected mini-batches to reduce the computation time per iteration. In this paper, we discuss the option of reducing the dimension of *H* via uniform sampling the rows to form a coreset 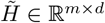 where *m* ≤ *n*. By reducing the sample size from *n* to *m*, we lessen the computational effort. As will be shown in our Theorem 1, the logistic loss is preserved for all *β*, and this suggests that 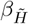 is a good approximation for *β*_*opt*_, where 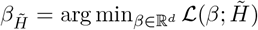. In addition, for matrix *H* ∈ ℝ^*n*×*d*^, we define its rank *k* approximation as *H*^*′*^ ∈ ℝ^*n*×*d*^ with 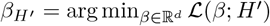.

Our first main theorem is to show that PCA and coreset preserve the logistic loss.

##### Theorem 1 (Main).

*Define H* ∈ ℝ^*n*×*d*^ *and its rank k approximation H*^*′*^ ∈ ℝ^*n*×*k*^. *For small classification complexity measure µ*(*H*^*′*^) *(see Definition 1), we can sample* 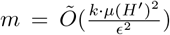 *rows from H to form its coreset* 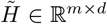. *After running logistic regression on* 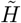 *to get* 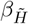, *we can use this parameter to predict the labels for H. The logistic loss* 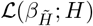 *will not deviate too much from* 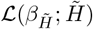, *which is a minimized value. More formally, with high probability*,

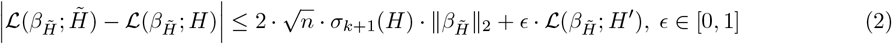

##### Proof Outline for Theorem 1

We bound the absolute difference in logistic loss between the original matrix *H* and its coreset *H*^*′*^ for the same parameter vector *β* in a straightforward way using the triangle inequality: We show the logistic loss is preserved between matrix *H* and its rank *k* approximation *H*^*′*^. Then we assert if *H*^*′*^ is *µ*-complex (i.e. *µ*(*H*^*′*^) is small), then we can construct a coreset matrix 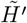 where the logistic loss is preserved. The absolute difference of the logistic loss between 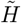 and 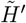 is also bounded analogously.

The full proof can be found in the Supplementary Section 3. The coreset construction technique (i.e. uniform balanced sampling) will be explained in detail in Section 2.2. We assume that the single-cell dataset *H* has finite samples *n* and *β*_*H*_*′* has bounded *ℓ*_2_ norm due to the regularization term. In addition, we assume the (*k* + 1)*th* largest singular value for *H* is very small. This assumption is reasonable because, after normalization and log-transformation, single-cell expression matrices typically exhibit a rapidly decaying singular value spectrum. Furthermore, the PCA step explicitly retains the top *k* dominant components, ensuring that the remaining singular values *σ*_*k*+1_(*H*), *σ*_*k*+2_(*H*), … are comparatively small.

#### Complexity Measure *µ*

The main caveat of Theorem 1 is that the size of the coreset depends on a somewhat opaque quantity, denoted *µ*(*H*^*′*^). Intuitively, *µ* captures the maximum imbalance between correctly classified vs. incorrectly classified points, that is how separable the dataset is. And it is known that there are datasets with high *µ* value that do not admit a small coreset [10]. Formally, *µ* is defined as the following.

##### Definition 1

([10, Definition 2. Classification Complexity Measure *µ*]). *For any X* ∈ ℝ^*n*×*d*^ *and y* ∈ {−1, 1}^*n*^, *let*

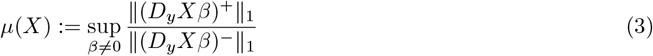

*where D*_*y*_ *is a diagonal matrix with y as its diagonal, and* (*D*_*y*_*Xβ*)^+^ *and* (*D*_*y*_*Xβ*)^−^ *denote the positive and the negative entries of D*_*y*_*Xβ, respectively*.

Our next theoretical result is to show that such *µ* must be small for single-cell data. More specifically, if we assume the rows of *H* are generated independently at random and has gone through PCA, we can show that *µ*(*H*^*′*^) must be near one, which coincides with our numerical experiments done in Section 3.1, due to a clever linear program to exactly compute *µ* [11]. We note that the original LP formulation in [11] contains a typographical error that was not consistently propagated through the subsequent derivations. We corrected and clarified this in Supplementary Section 4 (Lemma 3).

##### Theorem 2

(*µ* main (Informal)). *For label noise parameter ρ* ∈ (0, 1) *and same notations as before, with probability all but* ≤ 0.01, *we know µ*(*H*^*′*^) *is close to 1, that is*

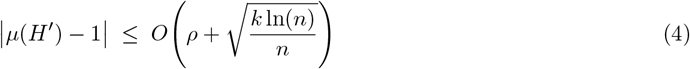

##### Proof Outline for Theorem 2

Our proof proceeds in three stages.

(1) We bound *µ* for a fixed *β* ∈ ℝ^*k*×1^ which can be formally stated as the following lemma.

#### Lemma 1

(Bound *µ* for A Fixed *β* ∈ ℝ^*k*×1^). *Define H* ∈ ℝ^*n*×*d*^ *and its rank k approximation H*^*′*^ ∈ ℝ^*n*×*k*^, *with a label noise parameter ρ* ∈ [0, 1] *and a constant < c* ∈ (0, 1] *where ρ < c. Then we can say, for a fixed β* ∈ ℝ^*k*×1^, *ϵ* ∈ (0, 1*/*2], *with probability all but* 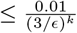,

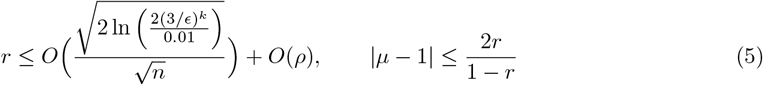

This involves (a) introducing the assumption on *H*^*′*^*β* under which the data yields a good vector *Y* (see Definition 1 in Supplementary Section 5), and we defer proving such assumption is valid to Step 3 of this outline; (b) applying Hoeffding’s inequality together with conditional independence of the labels to obtain a bound on *Q*, which is equivalent to the difference between numerator and denominator of (3) (See Lemma 4 in Supplementary Section 5); and (c) combining these ingredients to show that *µ* is bounded.

(2) Assuming Lemma 1, we extend the argument to hold for all *β* ∈ ℝ^*k*×1^, thereby proving the following lemma.

#### Lemma 2

(Bound *µ* for All *β* ∈ 𝕊^*k*−1^). *Using the same notations and assumptions as Lemma 1, with probability all but* ≤ 0.01, *for all unit β* ∈ 𝕊^*k*−1^,

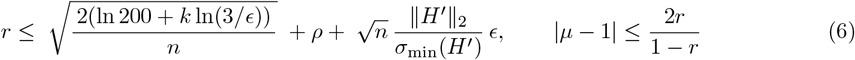

The key steps here are (a) strengthening the bound on *Q*, which is briefly introduced in step 1, by first using an *ϵ*-net of the unit sphere 𝕊^*k*−1^, 𝒩 of size |𝒩| ≤ (3*/ϵ*)^*k*^, which are roughly a set of points such that all points in 𝕊^*k*−1^ have *ϵ*-close point in 𝒩. We then use the union bound to give Lemma 1 type bound to all the points in 𝒩; (b) We show that *ϵ* deviation of *β* does not change the value of *r* much. Therefore our argument in (a) suffices to cover all *β* ∈ 𝕊^*k*−1^ (See Lemma 5 from Supplementary Section 5); (c) translating this uniform control of *Q* into a corresponding uniform bound on *µ*. Then, as a direct corollary from Lemma 2, we reach Theorem 2.

(3) Last but not least, we verify that the good vector *Y* assumption underlying our argument holds for the single-cell data under study (i.e. independent rows). An important technical ingredient is the near-uniform *ℓ*_2_ leverage score property.

The full proof can be found in Supplementary Section 5. Note that we can assume *σ*_min_(*H*^*′*^) here to be sufficiently large. If *σ*_min_(*H*^*′*^) were too small (i.e. close to zero), then the PCA pre-processing step would have discarded the corresponding low-variance directions. In other words, a small singular value indicates redundancy, and performing a more selective PCA would yield a reduced *H*^*′*^ with a larger *σ*_min_(*H*^*′*^). Recall we assumed *σ*_*k*+1_(*H*) to be small in Theorem 1. This remains consistent with the current claim. We can formally state the relationship as: for some positive threshold *t* > 0, *σ*_*k*+1_(*H*) ≤ *t* ≤ *σ*_min_(*H*^*′*^) = *σ*_*k*_(*H*^*′*^) = *σ*_*k*_(*H*).

### 2.2 Coreset construction

We construct coresets using a universal sampling approach that uniformly samples a fixed number of cells per label. A motivation for this strategy is provided below.

#### Uniform Sampling

We used uniform sampling to select rows into coresets instead of Lewis weights as stated in in Proposition 2 (Supplementary Section 4). In fact, after normalizing Lewis weights to lie between zero and one, the probabilities of selecting rows (i.e. cells) vary tightly around 1/*n* which is used in uniform sampling, see Figure 1. In our experiments, we observed nearly uniform Lewis weights for all datasets (data not shown).

**Figure 1.**
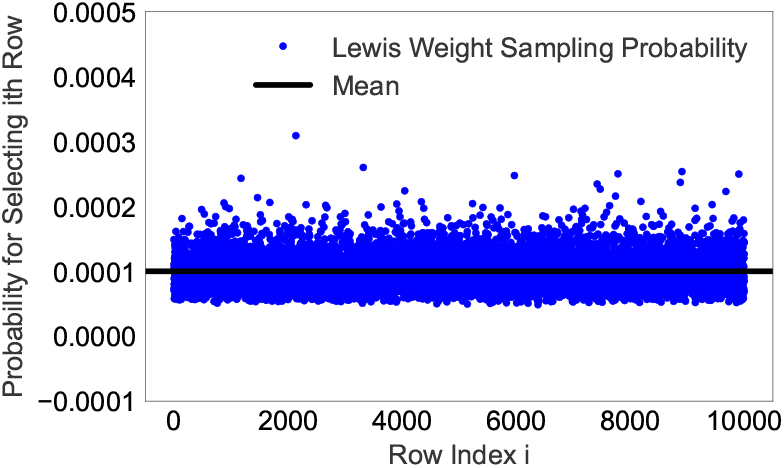
Lewis weights for 10k randomly selected cells from the hECA dataset.

#### Balanced Sampling

Recall that Theorem 1 implies that sampling 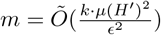 rows from the original matrix *H* is sufficient to construct a coreset 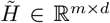. Given Theorem 2, this sampling size *m* can be regarded as a constant value. Consequently, for a small constant error *ϵ*, the required coreset size reduces to Õ(1), where the tilde notation hides a logarithmic factor in *d* and *µ*(*H*^*′*^), and a log-log factor in *n* [8]. In our application, instead of solving the multi-class classification problem directly, we decompose it into binary classification problems, one for each cell type, using the one-versus-rest (OvR) strategy. Accordingly, the theoretical Õ(1) coreset size represents the constant number of rows required per label. To exploit this theoretical guarantee in practice, we employ balanced sampling, selecting an equal number of cells from each cell type to ensure a uniform representation within the coreset.

## 3 Results

We evaluate our coreset-based logistic regression method implemented in tool CoreLR on two tasks: (i) cell type classification from single-cell and single-nucleus RNA-seq data, and (ii) transfer of spatial niche labels inferred from spatial transcriptomics data. The goal of these experiments is not to exhaustively benchmark against existing methods, but rather to demonstrate the effectiveness of coreset-based acceleration of logistic regression for single-cell and spatial omics analysis.

For the cell type prediction task we selected five datasets. We included two datasets that were used in [12] to benchmark the single-cell language model scMulan. One contained 60,345 human cardiac nuclei from the left ventricle (LV) in 8 healthy controls [13] and one in which the authors in [12] sampled 200,000 cells from the human Ensemble Cell Atlas (hECA), which contains labeled human cells from 116 published datasets, covering 38 organs and 11 systems. Moreover, we benchmarked methods on a human skin atlas of 710,704 cells [14], on 504,278 nuclei from the cardiovascular (CV) system of the healthy Wistar rat [15], and on single-nuclei RNA-seq of 111,907 cells from the dorsal vagal complex (DVC) of mouse and rat [16]. Following the strategy in [7], all datasets were preprocessed by normalizing counts to 10,000 and log transformed after adding a pseudocount of 1. Gene expression was standardized to zero mean and unit variance. The hECA dataset contained 2,000 highly variable genes that the authors selected when benchmarking scMulan in [12].

### 3.1 Complexity measure *µ*

We implemented the linear program introduced in Lemma 3 (Supplementary Section 4) to compute the classification complexity measure *µ* using Gurobi. Because the linear program is defined for binary labels, whereas our single-cell datasets contain multiple classes, we computed *µ* for each cell type in a one-versus-rest setting and reported the averaged *µ* values in Table 1 for all five datasets using varying numbers of principal components. A *µ* value of ∞ indicates that the dataset is exactly separable, that is, a hyperplane exists that perfectly separates all points of one class from those of the other. Accordingly, a large *µ* value (and especially *µ* = ∞) reflects poor classification complexity, implying that the dataset is ill-suited for logistic regression. By contrast, the *µ* values obtained after PCA (Table 1) are relatively small and close to one, indicating favorable conditions for logistic regression. Theorem 2 further guarantees that all PCA-transformed single-cell datasets yield good *µ* values near 1.

**Table 1:** Complexity measure *µ* for varying number of principal components (#PCs) on uniformly sampled 10k rows for all five datasets.

| #PCs | 10 | 50 | 100 | 500 | 1000 |
| --- | --- | --- | --- | --- | --- |
| LV | 1.122 | 1.100 | 1.082 | 1.041 | 1.030 |
| DVC | 1.128 | 1.082 | 1.060 | 1.022 | 1.015 |
| hECA | 1.001 | 1.001 | 1.001 | 1.000 | 1.000 |
| CV | 1.041 | 1.025 | 1.017 | 1.006 | 1.004 |
| Skin | 1.061 | 1.040 | 1.029 | 1.012 | 1.008 |

As shown in Table 1, *µ* decreases as the number of principle components *k* increases, suggesting that the worst-case bound established in Theorem 2 may be loose in practical regimes. Developing a refined analysis that narrows this gap between theoretical and empirical behavior represents a promising direction for future work. Importantly, the observation that *µ* approaches 1 with increasing *k* should not be misinterpreted as implying *µ* = 1 in the original feature space. On the contrary, in the absence of PCA where no theoretical guarantee applies, *µ* can diverge and even become unbounded, reflecting instability in the original high-dimensional representation. The corresponding experimental results from running the linear program without enforcing binary labels are provided in Supplementary Section 2, where the *µ* values for untransformed data could reach the extreme case of ∞.

### 3.2 CoreLR accurately and efficiently classifies cell types

We evaluated logistic loss and balanced classification accuracy when constructing coresets of varying sizes with and without prior PCA transformation. We selected 100 PCs on all datasets. Balanced accuracy accounts for imbalanced cell type compositions and is more sensitive to rare cell types. The observed coreset sizes are typically smaller than the target coreset ratios whenever the target ratio exceeds the observed frequency of at least one cell type, which limits the number of samples available for balanced sampling (see Supplementary Table 1). We compared our coreset-based approach CoreLR to CellTypist, which follows an alternative strategy to speeding-up training a (*ℓ*_2_-regularized) logistic regression model. It splits training data into mini-batches (1000 by default) and trains the model by stochastic gradient descent which led to an enormous speed-up compared to traditional logistic regression in [17]. Ground truth labels were obtained from the original publications.

We split the data into 80% for training and 20% for testing. In CoreLR we trained a *ℓ*_2_-regularized logistic regression model using function LogisticRegression of the scikit-learn pwhich are roughly a set ofackage. We performed 3-fold cross validation on a coreset of size 10% instead of the entire dataset to further speed-up our approach. The PCA-based version of CoreLR was evaluated on the original 20% test set by projecting the reduced coefficients back to the full *d*-dimensional feature space using the principal component loadings.

#### 3.2.1 Coresets provide near-identical loss

The theoretic guarantee on the coreset-based logistic loss provided by Theorem 1 is supported by experiments on all five datasets. In Figure 2, we observe near-identical loss on the entire matrix (orange) and its coreset (green) after rank *k* approximation (here PCA; dotted lines) for all five datasets. Even when omitting the PCA transformation, orange and green solid lines show only a small gap on datasets hECA, Skin and CV which further decreases with increasing coreset size. The gaps between the solid green and orange lines are slightly larger for the DVC and LV data, possibly due to an underrepresentation of rare cell types that occur at lower frequencies than the target sampling rate. Across all datasets and across all coreset sizes, the theoretic guarantees on the training loss yielded a more accurate solution, that is, a smaller logistic loss on the test data (blue lines) compared to the model trained by CellTypist (red line).

**Figure 2.**
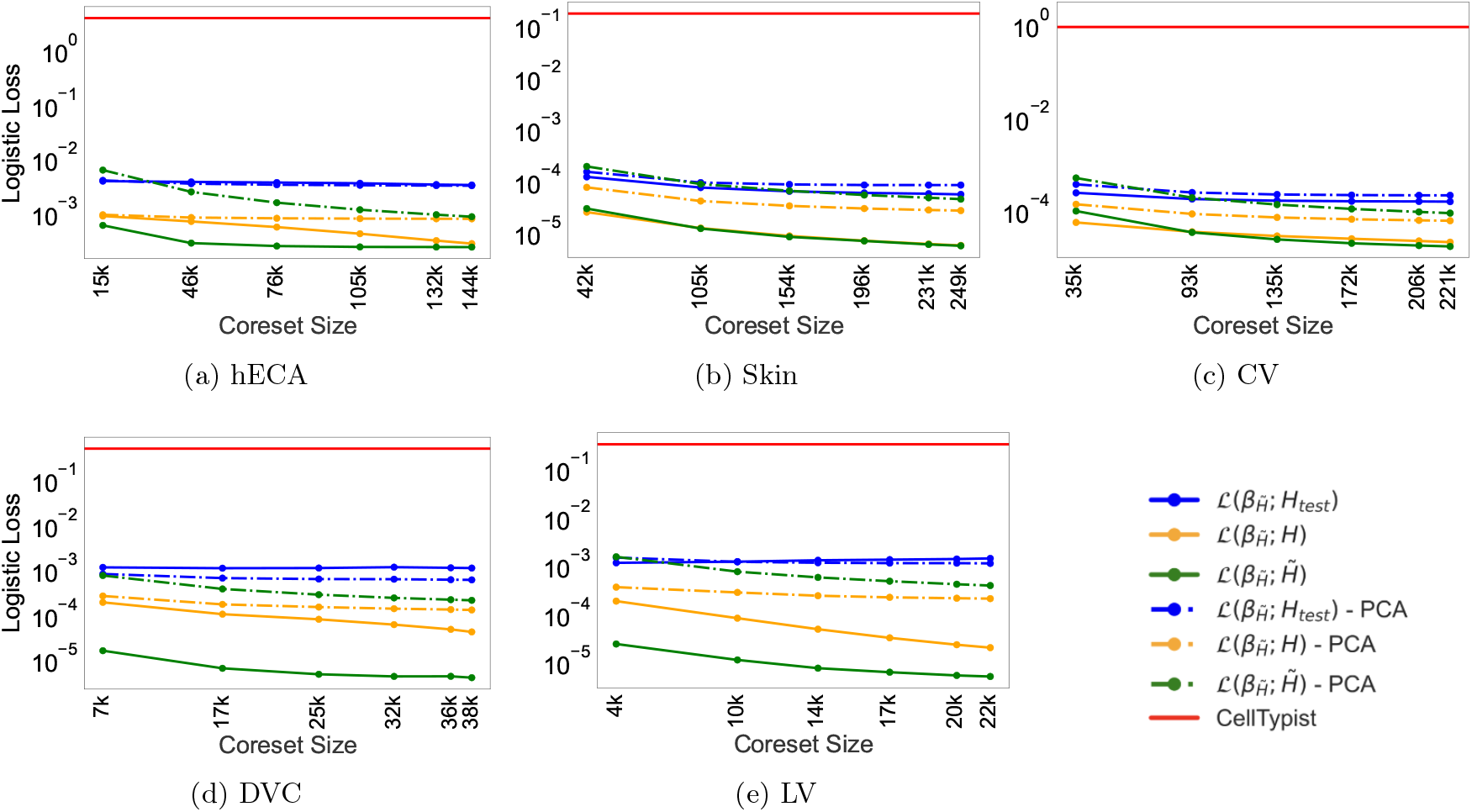
Logistic loss (without regularization term) in log scale as a function of observed coreset size. The observed coreset sizes shown on the x-axis correspond to target coreset ratios 0.1, 0.3, 0.5, 0.7, 0.9, and 1.0. Logistic loss is shown for the original matrix *H* (orange), its coreset (green), and on the test data (blue), with (dotted line) and without (solid line) prior PCA transformation, and compared to CellTypist’s loss (red) on the same test data.

#### 3.2.2 Coresets yield accurate cell type classification

Although classification accuracy is not directly controlled by our theory and depends on specific properties of the dataset such as the relative abundance of cell types, gene expression similarity between cell (sub-)types, and heterogeneity of cell states, Figure 3 demonstrates that a theoretically guaranteed small logistic loss results in accurate cell type classification in practice and outperforms a competing cell type annotation approach that utilizes the entire dataset. The slight decreasing trend observed in Figure 3 (b)-(e) as the coreset size increases might be attributed to two factors: (1) potential overfitting with larger coresets, and (2) reduced per-class accuracy for cell types that are underrepresented in the sketch due to the target sampling rate surpassing their observed frequency. For the experiments shown in Figure 3, we additionally ran CellTypist with its built-in “top genes” parameter set to 100. In this configuration, CellTypist selects the top 100 genes for each label based on their absolute regression coefficients; the final feature set used for classification is the union of these selected genes across all labels. This additional feature selection step had a negligible effect on the balanced accuracy of CellTypist (Figure 3).

**Figure 3.**
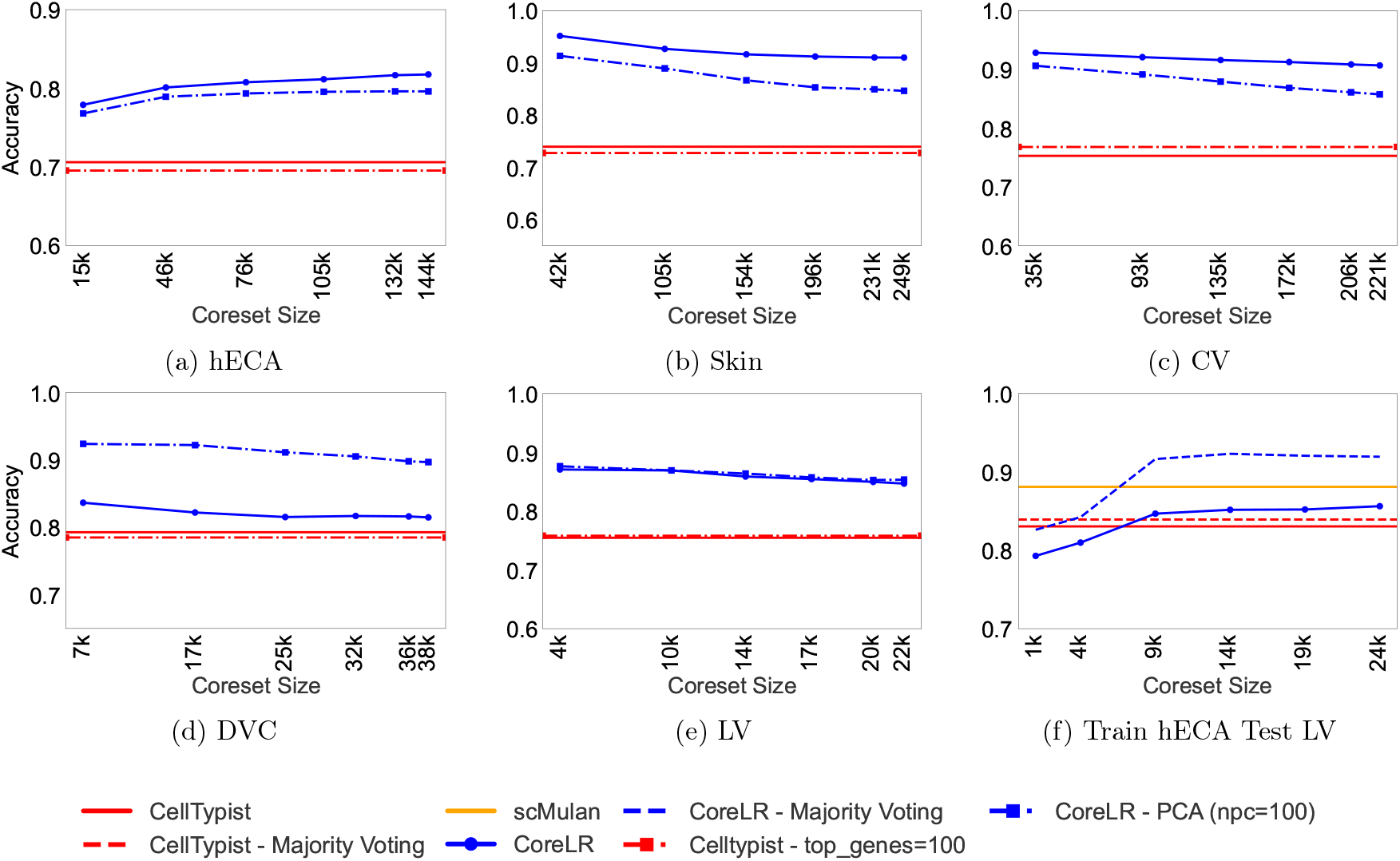
Accuracy as a function of observed coreset size. (a)-(e): Balanced accuracy of CoreLR (blue) with and without PCA (dotted and solid lines, respectively) and of CellTypist (red) with and without selecting the top 100 genes (dotted and solid lines, respectively) achieved on 20% test data. The observed coreset sizes shown on the x-axis correspond to target coreset ratios 0.1, 0.3, 0.5, 0.7, 0.9, and 1.0. (f): Accuracy of CoreLR (blue), CellTypist (red), and scMulan (orange) when training on hECA and testing on LV. Dashed blue and dashed red lines indicate additional majority voting for CoreLR and CellTypist, respectively. The x-axis represents observed coreset sizes corresponding to target ratios of 0.005, 0.025, 0.05, 0.075, 0.1, and 0.125 of in total 200,000 cells, shown from left to right.

Finally, we evaluated the generalization ability of the different models by training them on the hECA dataset and testing them on the LV data. This is the zero-shot setting without fine-tuning in which the cell type annotation performance of scMulan was benchmarked in [12]. To harmonize cell type labels between datasets, we used the label mapping provided by the authors in [12]. In addition, we followed the evaluation scheme in [12] by not considering the identification of a true cell type label as one of its subtypes as a false positive prediction. For testing on LV data, we selected the same 2,000 genes that were used when training on the hECA dataset.

Even in this zero-shot setting, CoreLR achieved results comparable to scMulan (Figure 3f). With a simple majority voting scheme similar to that applied by CellTypist in [7], CoreLR predicted cell types with an overall accuracy of 91.6%—achieved using only 5% of the data. In contrast, scMulan’s complex language model reached 88.1% accuracy when trained on the entire set of 10 million cells, while CellTypist achieved 83.9% overall accuracy after majority voting.

#### 3.2.3 Running times

Besides theoretical guarantees on the logistic loss, the main advantage of the coreset-based approach is its efficiency. The experiments in this work were run on Intel Xeon Gold 6226R CPU with 8 cores. Figure 4 left compares the running times of CellTypist and CoreLR for performing cross-validation. On the hECA dataset, CellTypist required about one order of magnitude more computation time than CoreLR for 3-fold cross-validation. Figure 4 right shows the computation times for all five datasets, using the optimal regularization parameter C selected during cross-validation. As expected, CoreLR with PCA achieves the lowest overall computation time, even when including both the PCA transformation and model training. In contrast, CellTypist, with or without the selection of the top 100 genes, consistently incurs the highest computational cost across all datasets. On the LV dataset, for example, CellTypist requires approximately 1.5 orders of magnitude more time than CoreLR with PCA.

**Figure 4.**
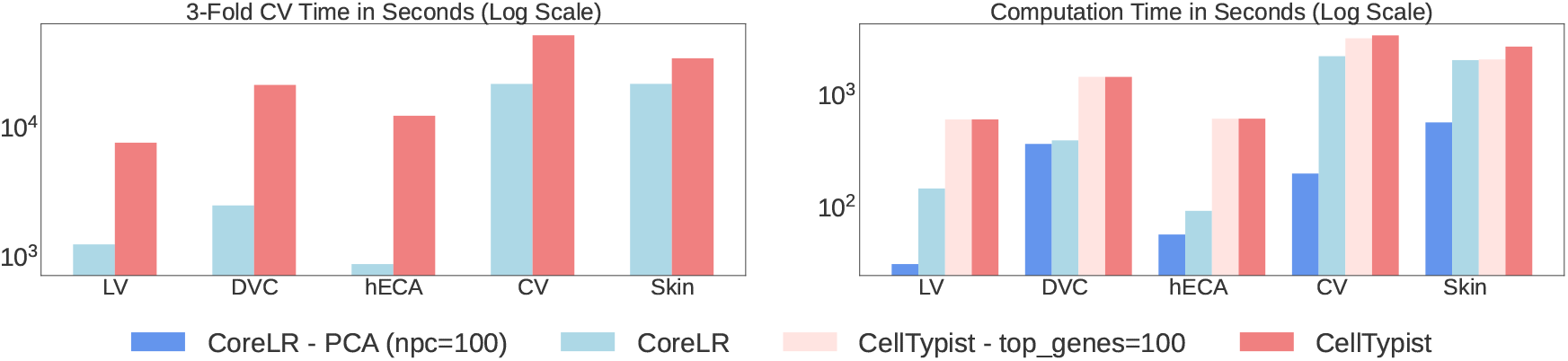
Total runtime in seconds (log scale) for (left) 3-fold cross-validation on the negative logistic loss using an intended coreset ratio of 0.1, and (right) training CoreLR (intended coreset ratio = 0.1) compared with CellTypist, with and without selecting the top 100 genes. All runtimes are reported with and without PCA preprocessing, including PCA computation time when applied.

Remarkably, to reach a level of accuracy that is comparable to that of scMulan (Figure 3f), our approach required only 56 seconds to train on just 5% of cells, compared to 192 hours scMulan needed for training on 10 million cells on four Nvidia-A800 GPUs. For comparison, the CellTypist approach needed 633 seconds for training. Furthermore, annotating cell types in the LV dataset based on the pre-trained scMulan model exceeded 9 hours of computation time, which the authors suggest can be reduced substantially using GPUs [12]. In contrast, predicting cell types took logistic regression based methods around one second.

### 3.3 Coresets allow accurate transfer of spatial niches across tissue sections

Besides predicting and transferring cell types across large datasets, we hypothesized that logistic regression could also support the transfer of tissue niches, i.e. spatially colocalized cell communities that coordinate biological function. Reliable niche transfer enables consistent biological interpretation and comparative analysis across diverse spatial transcriptomics platforms and moves us closer to a unified atlas of spatial niches. NicheCompass was recently introduced to identify biologically meaningful niches based on cellular communication pathways and gene program activity [18]. Its graph-based formulation encodes gene expression in the context of each cell’s local microenvironment and promotes interpretability by aligning embedding dimensions with signaling events.

Here, we asked whether niche labels identified by NicheCompass could be transferred from one dataset (training) to another (testing) without relying on prior pathway knowledge or learning spatial programs de novo. To this end, we applied our CoreLR model to the two largest datasets analyzed in the original NicheCompass study: the 313-probe Xenium human breast cancer dataset [19] comprising 286, 523 cells across two replicates, and the Nanostring CosMx human non-small cell lung cancer (NSCLC) dataset [20] from two donors (donor 5, adenocarcinoma, 76, 536 cells; donor 6, squamous cell carcinoma, 94, 673 cells.).

To make use of spatial neighborhood information in our approach, we first applied an approximated spectral graph transformation [21][22] to represent each spatial dataset in a lower-dimensional graph space.

For each dataset, CoreLR was trained on this representation of one sample and predicted niche labels for the held-out sample. The resulting labels were further refined based on the labels of the *k*-nearest neighbors. In addition to CellTypist, we compared performance to a graph isomorphism network (GIN) and a baseline random forest (RF) classifier which were recently shown to be able to accurately classify tissue phenotypes based on spatial expression patterns [18]. All methods were provided the identical low-dimensional graph embedding. Ground truth labels were inferred by NicheCompass.

Niche labels transferred by CoreLR across replicates and donors were consistent with the integrated embedding of samples by NicheCompass and matched tissue architecture (Figure 5). Compared to competing methods, CoreLR transferred labels with a substantially higher accuracy to all four samples (Figure 6, left). This quantitative improvement can be visually confirmed in the tissue sections shown in Figures 7 and 8. For example, CoreLR was the only method that correctly transferred fibroblast-endothelial cells from replicate 2 to replicate 1 in the Xenuim breast cancer dataset and plasma cells from donor 6 to donor 5 in the NanoString CosMx NSCLC dataset. At the same time, CoreLR achieved orders of magnitude faster training times (Figure 6, right), suggesting its suitability for large-scale spatial atlas construction.

**Figure 5.**
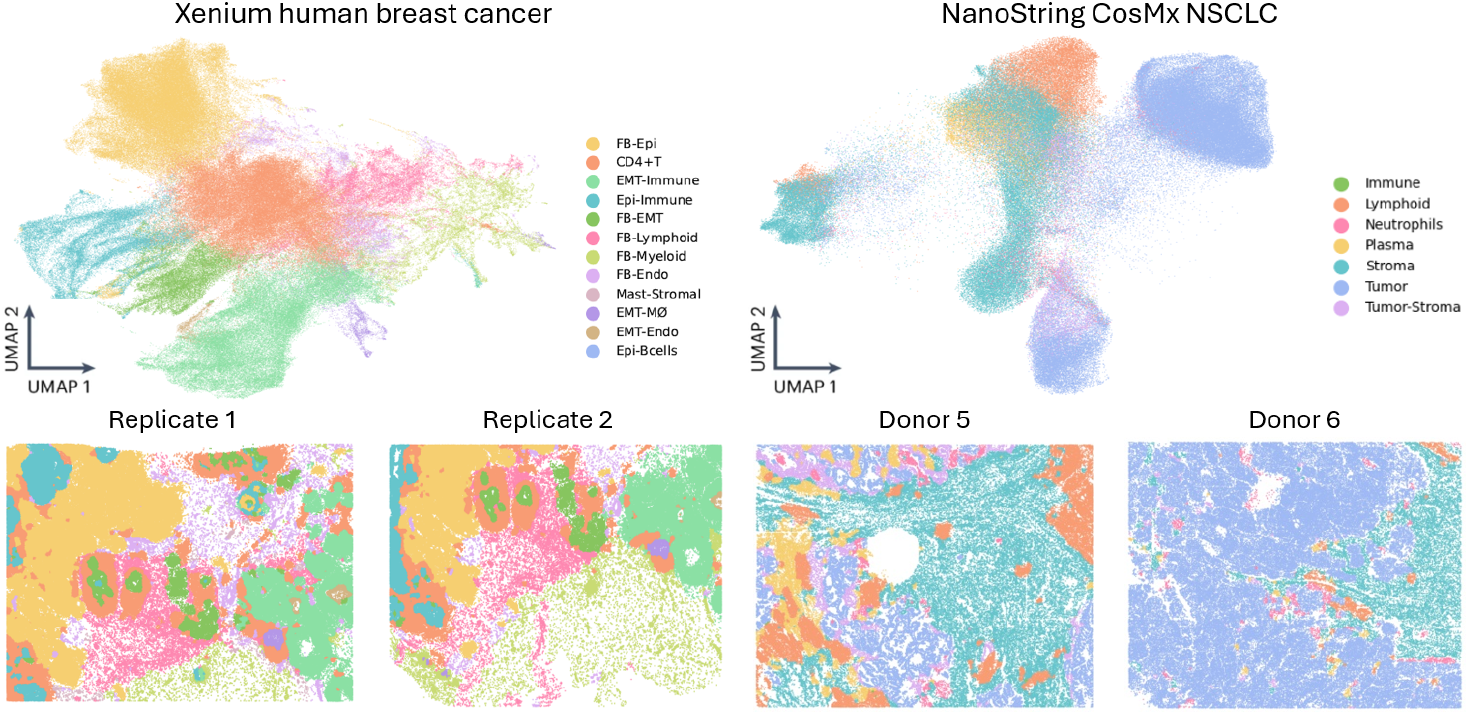
Top: UMAP visualization of the NicheCompass embedding of two replicates of the Xenium human breast cancer dataset (left) and two samples (donors 5 and 6) from the NanoString CosMx human NSCLC dataset (right). Colors indicate niches identified by CoreLR: FB, fibroblast; Epi, epithelial; Endo, endothelial; M, macrophage. Bottom: Corresponding tissue sections for the two replicates (left) and donors 5 and 6 (right), shown using the same niche color scheme as in the UMAP.

**Figure 6.**
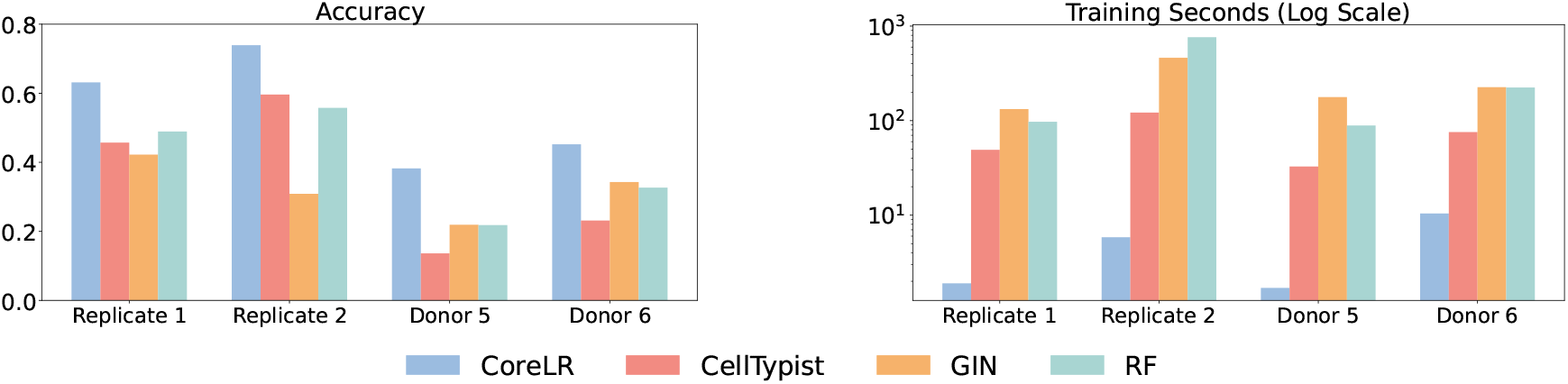
Balanced accuracy (left) and training time in seconds (right) for CoreLR, CellTypist, graph isomorphism network (GIN), and Random Forest (RF) on two replicates of the Xenium human breast cancer dataset and two samples from the NanoString CosMx human NSCLC dataset. Shown are test accuracies for replicate 1 (replicate 2) when training on replicate 2 (replicate 1), and analogously for donors 5 and 6.

**Figure 7.**
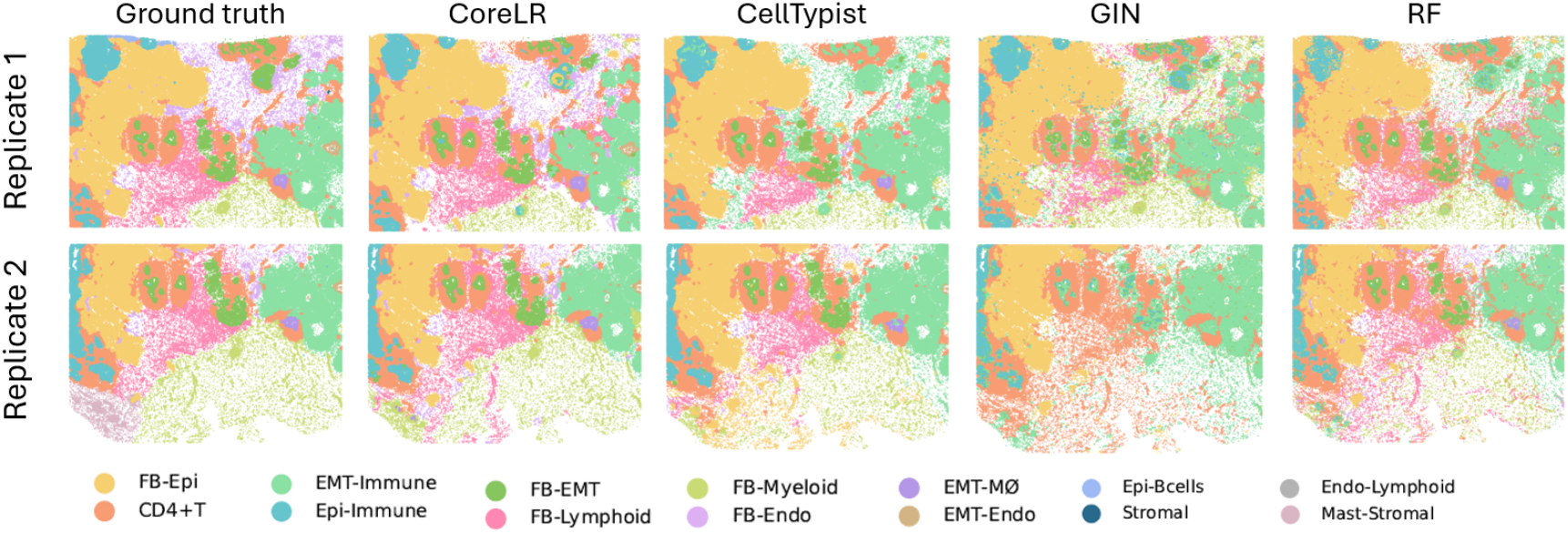
Tissue sections from two replicates of the human breast cancer dataset, shown with ground truth labels (first column) and with niche labels transferred by CoreLR, CellTypist, graph isomorphism network (GIN), and Random Forest (RF). Labels were transferred from replicate 2 to replicate 1 (top row) and from replicate 1 to replicate 2 (bottom row). FB, fibroblast; Epi, epithelial; Endo, endothelial; M, macrophage.

**Figure 8.**
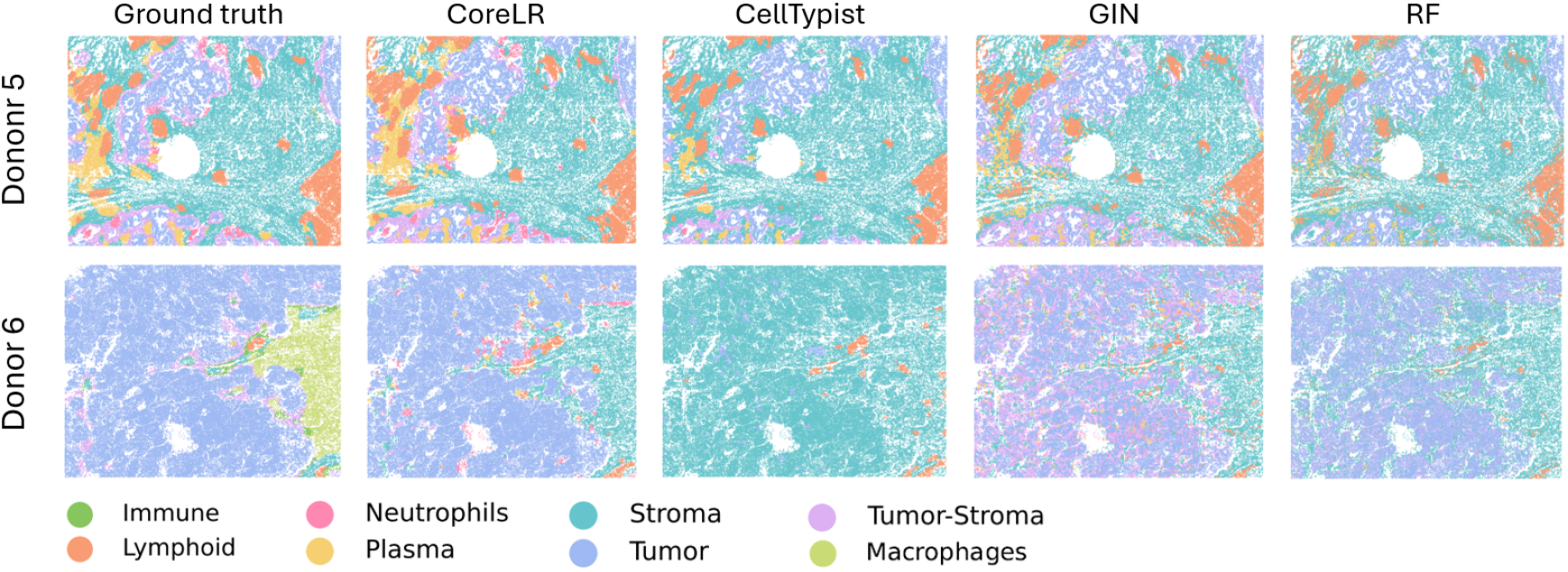
Tissue sections from two donors of the NanoString CosMx human NSCLC dataset, shown with ground truth labels (first column) and with niche labels transferred by CoreLR, CellTypist, graph isomorphism network (GIN), and Random Forest (RF). Labels were transferred from donor 6 to donor 5 (top row) and from donor 5 to donor 6 (bottom row).

## 4 Conclusion

In this work, we introduced CoreLR: a coreset-based approach for accelerating logistic regression with theoretical guarantees on the preservation of logistic loss. Our method enables efficient training on large-scale single-cell and spatial omics datasets while maintaining high predictive accuracy. We proved theoretically, and confirmed empirically, that single-cell datasets exhibit low classification complexity after PCA transformation, a dimensionality reduction step that is routinely applied in scRNA-seq analysis. Across multiple cell atlases and spatial transcriptomics datasets, we demonstrated that these theoretical guarantees translate into accurate and scalable transfer of both cell type and spatial niche labels, outperforming more complex models. Moreover, training logistic regression models on coresets is sufficiently fast to enable the transfer of cell type labels from large-scale reference atlases, such as immune or tissue-specific atlases, to individual single-cell studies, as well as the transfer of spatial labels from spatial omics datasets that currently comprise hundreds of images and are expected to expand substantially in the future.

Even though we sample rows (cells) uniformly at random, our theoretical guarantees are stronger than previous results based on uniform sampling [23]. As noted in our Lewis weights calculation, the weights are near uniform. Our conjecture is that if rows are i.i.d. samples (like cells in single-cell omics experiments), Lewis weights must be within some constant factor of the uniform distribution with high probability.

We note that, despite the proposed subsampling of cells for label transfer, single-cell and spatial omics studies still benefit greatly from profiling large numbers of cells. For instance, clustering cells based on their overall transcriptional similarity can reveal rare cell types or previously unknown cellular states, provided that a sufficiently large number of such cells are captured and sequenced. Indeed, more sophisticated sketching methods, such as Sphetcher [24], explicitly leverage large datasets by sampling cells that accurately represent the transcriptional heterogeneity of the entire population. In the supervised setting considered in this work, cell types or spatial niches are assumed to be known during training, enabling a balanced sampling strategy that ensures adequate representation of rare cell types within the coreset.

In our benchmark, scMulan, pre-trained on 10 million cells, achieved slightly inferior results compared to logistic regression combined with a simple majority voting scheme, even in the more challenging zero-shot setting. This finding aligns with previous research [2]. Nevertheless, through fine-tuning, foundation models provide powerful tools for various other types of downstream analyses of single-cell datasets, and their capabilities will continue to improve as the size of the training datasets increases.

In this work, we focused on the predictive analysis of single-cell and spatial omics data. However, logistic regression is also widely used in other analytical contexts, such as association analyses including cell type proximity analysis, and has shown particularly strong performance in marker gene selection [1]. In future work, we plan to investigate the applicability of similar coreset-based strategies to these and other types of analyses. The source code of CoreLR, the linear program for computing *µ*, approximating Lewis weight, and reproducing the results of our experiments are available at https://github.com/danrongLi/coreset_based_log_reg. The datasets used in this study can be downloaded from the same repository.

## References

1. Pullin, J. M. & McCarthy, D. J. A comparison of marker gene selection methods for single-cell RNA sequencing data. en. Genome Biology 25, 56. issn: 1474-760X. (2024) (Feb. 2024).

2. Boiarsky, R., Singh, N., Buendia, A., Getz, G. & Sontag, D. A Deep Dive into Single-Cell RNA Sequencing Foundation Models en. Oct. 2023. (2025).

3. Vannan, A. et al. Spatial transcriptomics identifies molecular niche dysregulation associated with distal lung remodeling in pulmonary fibrosis. en. Nature Genetics 57, 647–658. issn: 1061-4036, 1546-1718. (2025) (Mar. 2025).

4. Ali, M., Richter, S., Ertürk, A., Fischer, D. S. & Theis, F. J. Graph neural networks learn emergent tissue properties from spatial molecular profiles. en. Nature Communications 16, 8419. issn: 2041-1723. (2025) (Sept. 2025).

5. Hao, M. et al. Large-scale foundation model on single-cell transcriptomics. en. Nature Methods 21, 1481–1491. issn: 1548-7091, 1548-7105. (2024) (Aug. 2024).

6. Cui, H. et al. scGPT: toward building a foundation model for single-cell multi-omics using generative AI. en. Nature Methods 21, 1470–1480. issn: 1548-7091, 1548-7105. (2024) (Aug. 2024).

7. Domínguez Conde, C. et al. Cross-tissue immune cell analysis reveals tissue-specific features in humans. en. Science 376, eabl5197. issn: 0036-8075, 1095-9203. (2024) (May 2022).

8. Mai, T., Musco, C. N. & Rao, A. Coresets for Classification – Simplified and Strengthened en. In Advances in Neural Information Processing Systems (Nov. 2021). (2024).

9. Daliri, M., Freire, J., Li, D. & Musco, C. Matrix Product Sketching via Coordinated Sampling 2501.17836 [cs]. Jan. 2025. (2025).

10. Munteanu, A., Schwiegelshohn, C., Sohler, C. & Woodruff, D. On Coresets for Logistic Regression in Advances in Neural Information Processing Systems 31 (Curran Associates, Inc., 2018). (2024).

11. Dexter, G., Khanna, R., Raheel, J. & Drineas, P. Feature Space Sketching for Logistic Regression 2303.14284 [cs, stat]. Mar. 2023. (2024).

12. Bian, H. et al. scMulan: a multitask generative pre-trained language model for single-cell analysis en. Jan. 2024. (2024).

13. Simonson, B. et al. Single-nucleus RNA sequencing in ischemic cardiomyopathy reveals common transcriptional profile underlying end-stage heart failure. en. Cell Reports 42, 112086. issn: 22111247. (2024) (Feb. 2023).

14. Fiskin, E. et al. Multi-modal skin atlas identifies a multicellular immune-stromal community associated with altered cornification and specific T cell expansion in atopic dermatitis en. Oct. 2023. (2025).

15. Portal, B. I. S. C. Transcriptional profile of the rat cardiovascular system at single cell resolution

16. Hes, C. et al. A unified rodent atlas reveals the cellular complexity and evolutionary divergence of the dorsal vagal complex en. Sept. 2024. (2025).

17. Cox, D. R. The Regression Analysis of Binary Sequences. en. Journal of the Royal Statistical Society Series B: Statistical Methodology 20, 215–232. issn: 1369-7412, 1467-9868. (2024) (July 1958).

18. Birk, S. et al. Quantitative characterization of cell niches in spatially resolved omics data. en. Nature Genetics 57, 897–909. issn: 1061-4036, 1546-1718. (2025) (Apr. 2025).

19. Janesick, A. et al. High resolution mapping of the tumor microenvironment using integrated single-cell, spatial and in situ analysis. en. Nature Communications 14, 8353. issn: 2041-1723. (2025) (Dec. 2023).

20. He, S. et al. High-plex imaging of RNA and proteins at subcellular resolution in fixed tissue by spatial molecular imaging. en. Nature Biotechnology 40, 1794–1806. issn: 1087-0156, 1546-1696. (2025) (Dec. 2022).

21. Defferrard, M., Bresson, X. & Vandergheynst, P. Convolutional neural networks on graphs with fast localized spectral filtering in Proceedings of the 30th International Conference on Neural Information Processing Systems (Curran Associates Inc., 2016), 3844–3852. isbn: 978-1-5108-3881-9.

22. Zhu, H. & Koniusz, P. Simple Spectral Graph Convolution in International Conference on Learning Representations (2021).

23. Curtin, R. R., Im, S., Moseley, B., Pruhs, K. & Samadian, A. On Coresets for Regularized Loss Minimization Version Number: 2. 2019. (2024).

24. Do, V. H., Elbassioni, K. & Canzar, S. Sphetcher: Spherical Thresholding Improves Sketching of Single-Cell Transcriptomic Heterogeneity. en. iScience 23, 101126. issn: 25890042. (2024) (June 2020).

