## Supplementary for "Coreset-based logistic regression for atlas-scale single-cell and spatial omics analyses"

#### 1 Intended vs Observed Coreset Ratio

| Intended Ratio | | 0.1 | 0.3 | 0.5 | 0.7 | 0.9 | 1.0 |
|---|---|---|---|---|---|---|---|
| Observed Ratio | hECA | 0.10 | 0.29 | 0.48 | 0.66 | 0.83 | 0.91 |
| | Skin | 0.08 | 0.19 | 0.27 | 0.34 | 0.41 | 0.44 |
| | CV | 0.09 | 0.23 | 0.34 | 0.43 | 0.51 | 0.55 |
| | DVC | 0.09 | 0.22 | 0.32 | 0.40 | 0.46 | 0.49 |
| | LV | 0.09 | 0.21 | 0.29 | 0.36 | 0.43 | 0.46 |

Table 1: Comparison of intended and observed coreset ratios on all five datasets.

#### 2 $\mu$ Values Without Binary Label

| #PCs | 10 | 100 | 1000 | no PCA |
|---|---|---|---|---|
| LV | 3.75 | 3.89 | 4.54 | $\infty$ |
| DVC | 3.21 | 3.37 | 3.80 | $\infty$ |
| hECA | 1.57 | 3.76 | 7.57 | $\infty$ |
| CV | 3.06 | 6.26 | 15.63 | $\infty$ |
| Skin | 4.21 | 5.60 | 6.15 | $\infty$ |

Table 2: Complexity measure $\mu$ for varying numbers of principal components (#PCs) on uniformly sampled 10k rows for all five datasets without enforcing binary labels.

Without enforcing binary labels for our LP implementation, we encoded class labels using scikit-learn's LabelEncoder (i.e., integer codes $y \in \mathbb{Z}_{\geq 0}$). This differs from the $\{\pm 1\}$ assumption in the definition of $\mu$: instead of only flipping row signs, $D_y$ rescales rows, and zeros those with $y = 0$. Consequently, $Q = \mathbf{1}^\top D_y H' \beta = y^\top H' \beta$ need not vanish even when $H'$ is column centered. This leads to non-negligible $r = |Q|/P$

[*]Corresponding authors

and empirical $\mu = \frac{1+r}{1-r} > 1$, matching the first three columns of Table 2. Without PCA, the $\mu$ values approach $\infty$ for all five datasets as shown in the last column.

### 3 Proof of Theorem 1

*Proof.* We first propose that a low-rank approximation of the original data preserves the logistic loss for the same regression coefficients.

**Proposition 1** (Logistic Loss Preservation after Rank $k$ Approximation)**.** *Assume the same notations as described previously. Recall $H' \in \mathbb{R}^{n\times d}$ represents the rank $k$ approximation for $H$. Then for all $\beta \in \mathbb{R}^d$,*

$$|\mathcal{L}(\beta; H) - \mathcal{L}(\beta; H')| \leq \sqrt{n} \cdot \sigma_{k+1}(H) \cdot \|\beta\|_2 \tag{1}$$

*Proof.* We will combine 2 straightforward lemma to conclude the proof.

**Lemma 1** ([1, Theorem 3])**.** *If $X, X' \in \mathbb{R}^{n\times d}$, for all $\beta \in \mathbb{R}^d$,*

$$|\mathcal{L}(\beta; X) - \mathcal{L}(\beta; X')| \leq \sqrt{n}\|X - X'\|_2\|\beta\|_2 \tag{2}$$

The right-hand-side of (2) can be bounded by the $(k+1)th$ largest singular value of the original matrix with the following standard lemma:

**Lemma 2** ([2, Eckart Young Mirsky Theorem])**.** *Suppose matrix $A'$ is the rank $k$ approximation for matrix $A \in \mathbb{R}^{n\times d}$. Then we have*

$$\|A - A'\|_2 = \sigma_{k+1}(A) \tag{3}$$

*where $\sigma_{k+1}(A)$ represents the $(k+1)th$ largest singular value for matrix $A$.*

With Lemma 1 and Lemma 2, we can show that under the condition that $H$ has a good rank $k$ approximation $H'$ (i.e. $\sigma_{k+1}(H)$ is small), implying that the logistic loss between $H'$ and $H$ differs by a small value, and this holds true for all regression coefficients $\beta \in \mathbb{R}^d$.

From Lemma 2, we get

$$\|H - H'\|_2 = \sigma_{k+1}(H) \tag{4}$$

while Lemma 1 gives

$$|\mathcal{L}(\beta; H) - \mathcal{L}(\beta; H')| \leq \sqrt{n}\|H - H'\|_2\|\beta\|_2 \tag{5}$$

By combining (4) and (5), we conclude the proof. □

We will now state Proposition 2 which argues that the coreset $\tilde{H}$ generated under the $\ell_1$ Lewis weight sampling (see [3]) preserves the logistic loss for the same regression coefficients.

**Proposition 2** ([4] Corollary 9 restated)**.** *Assume the same annotations as described before. Using the $\ell_1$ Lewis weight, suppose we sample $m = \tilde{O}(\frac{k\cdot\mu(H)^2}{\epsilon^2})$ rows from matrix $H \in \mathbb{R}^{n\times k}$ to construct coreset $\tilde{H} \in \mathbb{R}^{m\times k}$, where $\epsilon \in [0,1]$ and $\mu(H)$ is the complexity measure. Then with high probability, for all $\beta \in \mathbb{R}^k$, we have:*

$$\left|\mathcal{L}(\beta; \tilde{H}) - \mathcal{L}(\beta; H)\right| \leq \epsilon \cdot \mathcal{L}(\beta; H) \tag{6}$$

Proposition 2 suggests that after creating the coreset $\tilde{H}$, we can calculate its logistic regression coefficient $\beta_{\tilde{H}}$. Then with high probability:

$$\left|\mathcal{L}(\beta_{\tilde{H}}; \tilde{H}) - \mathcal{L}(\beta_{\tilde{H}}; H)\right| \leq \epsilon \cdot \mathcal{L}(\beta_{\tilde{H}}; H), \quad \epsilon \in [0,1] \tag{7}$$

Recall that Proposition 1 and Proposition 2 both have the guarantee on all $\beta \in \mathbb{R}^d$. By substituting $\beta_{\tilde{H}}$ into $\beta$ from (1) in Proposition 1, we have

$$|\mathcal{L}(\beta_{\tilde{H}}; H) - \mathcal{L}(\beta_{\tilde{H}}; H')| \leq \sqrt{n} \cdot \sigma_{k+1}(H) \cdot \|\beta_{\tilde{H}}\|_2 \tag{8}$$

By substituting $\beta_{\tilde{H}}$ into $\beta$ from (6) and using the rank $k$ approximation matrix $H'$, with high probability, we have

$$\left|\mathcal{L}(\beta_{\tilde{H}};\tilde{H}') - \mathcal{L}(\beta_{\tilde{H}};H')\right| \le \epsilon \cdot \mathcal{L}(\beta_{\tilde{H}};H'), \quad \epsilon \in [0,1] \tag{9}$$

By substituting $\beta_{\tilde{H}}$ and $\tilde{H}$ into (5), we get

$$\begin{aligned} |\mathcal{L}(\beta_{\tilde{H}};\tilde{H}) - \mathcal{L}(\beta_{\tilde{H}};\tilde{H}')| \le \sqrt{n} \cdot \|\tilde{H} - \tilde{H}'\|_2 \cdot \|\beta_{\tilde{H}}\|_2 &\le \sqrt{n} \cdot \|H - H'\|_2 \cdot \|\beta_{\tilde{H}}\|_2 \\ &\le \sqrt{n} \cdot \sigma_{k+1}(H) \cdot \|\beta_{\tilde{H}}\|_2 \end{aligned} \tag{10}$$

After combining (8), (9) and (10) with triangle inequality, we get with high probability,

$$|\mathcal{L}(\beta_{\tilde{H}};H) - \mathcal{L}(\beta_{\tilde{H}};\tilde{H})| \le 2 \cdot \sqrt{n} \cdot \sigma_{k+1}(H) \cdot \|\beta_{\tilde{H}}\|_2 + \epsilon \cdot \mathcal{L}(\beta_{\tilde{H}};H'),\ \epsilon \in [0,1] \tag{11}$$

which completes the proof of Theorem 1.

□

### 4 LP Formulation to Find $\mu$

Even though computing $\mu$ was initially believed to be intractible [5], only possible to approximate by a polynomial factor in polynomial time, the authors in [1] designed a simple linear program to compute $\mu(X)$ exactly.

**Lemma 3** ([1, Theorem 2.]). *If the complexity measure $\mu_y(X)$ of Definition 1 is finite, it can be computed exactly by solving a linear program with $2n$ variables and $4n$ constraints. The linear program is constructed as follows:*
*Define*

$$\beta^* = \arg\max_{\beta \neq 0} \frac{\|(D_y X\beta)^+\|_1}{\|(D_y X\beta)^-\|_1} \tag{12}$$

*where $\beta^* \neq \arg\min_\beta \mathcal{L}(\beta; X)$. Replace $D_y X\beta$ with vector $z \in \mathbb{R}^n$, where $z = z_+ - z_-$. Suppose $z_+ \in \mathbb{R}^n$ contains the absolute value of the positive elements in $z$, and $z_- \in \mathbb{R}^n$ contains the absolute value of the negative elements in $z$. Since we want to ensure $z \in$ Range($D_y X$), we will introduce $P_R \in \mathbb{R}^{n\times n}$ which is the orthogonal projection to Range($D_y X$).*

*Then we have objective function:*

$$z^* = \arg\max_{z\in\mathbb{R}^n} \mathbf{1}^T(z_+ - z_-) \tag{13}$$

*Subject to:*

$$\begin{aligned} \sum_{i=1}^{2n} [z_+, z_-]_i &\le C \\ (\mathbf{I} - P_R)(z_+ - z_-) &= \mathbf{0} \\ z_+,\ z_- &\ge \mathbf{0} \end{aligned} \tag{14}$$

*where $C \in \mathbb{R}$ is any positive constant, and we used $C = 1$ in our LP problem construction.*

*We can calculate $z^*$ as*

$$z^* = z_+^* - z_-^* \tag{15}$$

*Since $z^* = D_y X\beta^*$, we can calculate $\mu(x)$ as:*

$$\mu(X) = \frac{\|(D_y X\beta^*)^+\|_1}{\|(D_y X\beta^*)^-\|_1} = \frac{\|(z^*)^+\|_1}{\|(z^*)^-\|_1} \tag{16}$$

Note in (12), we corrected the calculation for $\beta^*$ from [1]. While the original formulation in [5] follows Definition 1, the authors in [1] flipped the denominator and the numerator but failed to switch from max to min in the definition of $\beta^*$, which we believe is a result of a typo. We have verified that the slightly modified argument (essentially analogous) to that of given in [1] indeed gives Lemma 3.

According to [4], the value of $\mu(H)$ for matrix $H$ can become significantly large if there exists some parameter vector $\beta \in \mathbb{R}^d$ that causes huge discrepancy between correctly classified and misclassified labels. If $H$ is a good candidate for logistic regression problems, we generally observe that $\mu(H)$ remains relatively small. [4] also suggests that if $\mu(H)$ is small, we can construct a coreset $\tilde{H} \in \mathbb{R}^{m\times d}$ where $m = \tilde{O}(\frac{d\cdot\mu(H)^2}{\epsilon^2})$, $\epsilon \in [0, 1]$ where $\epsilon$ is a relative error parameter.

We also remark that this is a **sufficient** condition for coreset not a necessary nor equivalent condition. As remarked in [5], it is possible to have a data set $H$ which permits a good small coreset while having a bad $\mu(H)$ value.

### 5 Proof of Theorem 2

#### 5.1 Preliminaries and Notations

Using similar notations as in [5] and assuming we use $k$ principal components for performing PCA on matrix $H \in \mathbb{R}^{n\times d}$ to get its rank $k$ approximation $H' \in \mathbb{R}^{n\times k}$, we state $D_y$ as a diagonal matrix with $y$ as its diagonal where $y \in \{-1, 1\}^n$ is the label vector. To simplify notations, let $Y^{(\beta)} = H'\beta$. From Definition 1, denote the numerator of (3) as $U$ and the denominator as $D$.

Then let us introduce the notation

$$Q^{(\beta)} := U^{(\beta)} - D^{(\beta)} = \sum_{i=1}^{n}\sum_{j=1}^{k}(D_yH')_{ij}\beta_j = \sum_{i=1}^{n} y_iY_i^{(\beta)} \tag{17}$$

$$P^{(\beta)} := U^{(\beta)} + D^{(\beta)} = \|D_y^{(\beta)}H'\beta\|_1 = \|D_yY^{(\beta)}\|_1 = \sum_{i=1}^{n}|y_iY_i^{(\beta)}| = \sum_{i=1}^{n}|Y_i^{(\beta)}| \tag{18}$$

With the above definition, we can then introduce a crucial ratio

$$r^{(\beta)} := \frac{|Q^{(\beta)}|}{P^{(\beta)}} \tag{19}$$

Observe that $r^{(\beta)} \le 1$. Then we can define (and write)

$$\mu^{(\beta)}(H') := \frac{P^{(\beta)} + Q^{(\beta)}}{P^{(\beta)} - Q^{(\beta)}} = \frac{1 + \frac{Q^{(\beta)}}{P^{(\beta)}}}{1 - \frac{Q^{(\beta)}}{P^{(\beta)}}}$$

Then we can rewrite (3) as

$$\mu(H') \; = \; \sup_{\beta\neq 0} \mu^{(\beta)}(H'). \tag{20}$$

Furthermore, for any fixed $\beta \in \mathbb{R}^{k\times 1}$ and $\beta \neq 0$, we get

$$\frac{1 - r^{(\beta)}}{1 + r^{(\beta)}} \le \mu^{(\beta)}(H') \le \frac{1 + r^{(\beta)}}{1 - r^{(\beta)}} \tag{21}$$

$$|\mu^{(\beta)}(H') - 1| \le \frac{2r^{(\beta)}}{1 - r^{(\beta)}} \tag{22}$$

Note that if $r^{(\beta)}$ is around 0, then $\mu^{(\beta)}$ must be near 1. But if $r^{(\beta)}$ is near 1, then $\mu^{(\beta)}$ can be unbounded. So our whole goal is to show that under the conditional independence assumptions on the labels as well as additional restrictions on the entries of $Y$, we can say that $r^{(\beta)}$ must be near 0 for any given $\beta$.

**Fact 1** ([6, Hoeffding's Inequality])**.** *Let $F_1, \ldots, F_n$ be independent random variables with each $F_i \in [k_i, l_i]$. Let*

$$\overline{\mu} \;=\; \sum_{i=1}^{n} \mathbb{E}[F_i] \tag{23}$$

*Then for any $t > 0$, the sum $F = \sum_{i=1}^{n} F_i$ satisfies*

$$\Pr\big(|F - \overline{\mu}| > t\big) \;\leq\; 2\exp\Big(-\frac{2t^2}{\sum_{i\in n}(l_i - k_i)^2}\Big) \tag{24}$$

#### 5.2 Assumption Definition

To link the bound on $Q^{(\beta)}$ to $\mu^{(\beta)}$, we would like to have

$$R^{(\beta)} := \frac{\|Y^{(\beta)}\|_1}{\|Y^{(\beta)}\|_2} = \Theta(\sqrt{n}) \tag{25}$$

Note that $R^{(\beta)} \leq \sqrt{n}$ due to Cauchy-Schwarz, but we do not have any guarantees on its lower bound. To give a lower bound on $R^{(\beta)}$, we introduce the following definition on good $Y^{(\beta)}$.

**Definition 1** (Good $Y^{(\beta)}$)**.** *With the notations as described above, suppose $G \subseteq [n]$, $|G| \geq \frac{n}{c}$ where $c \in \mathbb{Z}^+$. If for all $i \in G$, with some positive constant $t \in \mathbb{Z}^+$, we have the following constraint,*

$$\frac{|Y_i^{(\beta)}|}{\|Y^{\beta}\|_2} > \frac{t}{\sqrt{n}} \tag{26}$$

The following claim then shows that if Definition 1 is satisfied, indeed $R^{(\beta)} = \Theta(\sqrt{n})$.

**Claim 1.** *If $Y^{(\beta)}$ satisfies Definition 1, then*

$$\frac{\|Y^{(\beta)}\|_1}{\|Y^{(\beta)}\|_2} = \Theta(\sqrt{n}) \tag{27}$$

*Proof.* Since Cauchy–Schwarz trivially leads to $\frac{\|Y^{(\beta)}\|_1}{\|Y^{(\beta)}\|_2} \leq \sqrt{n}$, it suffices to show $\frac{\|Y^{(\beta)}\|_1}{\|Y^{(\beta)}\|_2} \geq \Omega(\sqrt{n})$. With the definition of $|Y_i^{(\beta)}|$, we get

$$\|Y^{(\beta)}\|_1 \geq \sum_{i\in G} |Y_i^{(\beta)}| \geq \frac{n}{c} t \cdot \frac{\|Y^{(\beta)}\|_2}{\sqrt{n}} = \frac{t}{c} \cdot \sqrt{n} \cdot \|Y^{(\beta)}\|_2 \tag{28}$$

Dividing both sides by $\|Y^{(\beta)}\|_2$, we have

$$\frac{\left\|Y^{(\beta)}\right\|_1}{\left\|Y^{(\beta)}\right\|_2} \geq \frac{t}{c}\sqrt{n} \geq \Omega(\sqrt{n}) \tag{29}$$

Thus we can safely conclude the proof.

□

#### 5.3 Bound $Q^{(\beta)}$ for A Fixed $\beta \in \mathbb{R}^{k\times 1}$

In order to bound the deviation of $\mu^{(\beta)}(H')$ in Lemma 1, the next crucial step is to bound $Q^{(\beta)}$ with Hoeffding's bound. Recall (17), since $y_i$ represents the label for the $i$th sample, and $H'_i$ represents the feature vector after PCA for the same $i$th sample, it implies that $y_i$ depends on $H'_i$ for $i \in \{1, 2, 3, \ldots, n\}$.

Then we can assume the following. Given a label $y$, there exists a distribution on the rows that we sample from independently at random. This implies the conditional independence assumption as shown below.

$$
\begin{aligned}
\mathbb{P}[y|H'] &= \mathbb{P}[y_1, \dots, y_n|H'] \\
&= \prod_{i=1}^{n} \mathbb{P}\big[y_i|y_{<i}, H'\big] \quad \text{(chain rule)} \\
&= \prod_{i=1}^{n} \mathbb{P}\big[y_i|H'\big] \quad \text{(cond. independence)} \\
&= \prod_{i=1}^{n} \mathbb{P}\big[y_i|H'_i\big] \quad \text{(row-wise dependence)}
\end{aligned} \tag{30}
$$

We will use this assumption along with the Hoeffding's inequality to prove the following lemma.

**Lemma 4** (Bound $Q^{(\beta)}$ for A Fixed $\beta \in \mathbb{R}^{k\times 1}$)**.** *Given a PCA transformed matrix $H' \in \mathbb{R}^{n\times k}$, $Q^{(\beta)}$ is defined as (17), then with probability all but $\leq \frac{0.01}{(3/\epsilon)^k}$ and label noise parameter $\rho \in [0, 1]$, we have*

$$
|Q^{(\beta)}| \leq \|Y^{(\beta)}\|_2 \Big(\sqrt{2 \ln\Big(\frac{2(3/\epsilon)^k}{0.01}\Big)} + \rho\sqrt{n}\Big) \tag{31}
$$

*Proof.* With the conditional independence assumption, given the matrix $H'$, for any $i \in [n]$, denote $s_i := \mathbb{E}[y_i|H'_i] \in [-1, 1]$. Then we can define the base term $B_i^{(\beta)} := s_i Y_i^{(\beta)}$ and $B^{(\beta)} := \sum_{i=1}^n B_i^{(\beta)}$. Intuitively, this is the expected value of $Q_i^{(\beta)}$ and $Q^{(\beta)}$ respectively, under fixed $\beta$ and conditioned on $H'$.

Observe that each summand is bounded: $|Q_i^{(\beta)} - B_i^{(\beta)}| = |(y_i - s_i)Y_i^{(\beta)}| \leq 2|Y_i^{(\beta)}|$ for $i \in [n]$. Since $\overline{\mu} = \sum_{i=1}^n \mathbb{E}[y_i - s_i|H'] = 0$, applying Fact 1, we can say with probability all but $\frac{0.01}{(3/\epsilon)^k}$,

$$
|Q^{(\beta)} - B^{(\beta)}| \leq \|Y^{(\beta)}\|_2 \sqrt{2 \ln\Big(\frac{2(3/\epsilon)^k}{0.01}\Big)} \tag{32}
$$

Recall that $H' \in \mathbb{R}^{n\times k}$ is the PCA transformed matrix. We achieve this via `sklearn.decompositio n.PCA` package, which by default, centers features by subtracting the sample mean. In other words, $H'$ is column centered. Therefore $\sum_{i=1}^n Y_i^{(\beta)} = \sum_{i=1}^n {H'}_i^\top \beta = \sum_{j=1}^k \beta_j \Big( \sum_{i=1}^n H'_{ij} \Big) = 0$.

Denote $\overline{s} = \frac{1}{n}\sum_{i=1}^n s_i$ and $d_i = s_i - \overline{s}$ for $i \in [n]$. Then we can write

$$
B^{(\beta)} = \sum_{i=1}^n s_i Y_i^{(\beta)} = \sum_{i=1}^n (d_i + \overline{s}) Y_i^{(\beta)} = \sum_{i=1}^n d_i Y_i^{(\beta)} + \overline{s} \underbrace{\sum_{i=1}^n Y_i^{(\beta)}}_{=0} = d^\top Y^{(\beta)} \leq \|d\|_2 \|Y^{(\beta)}\|_2 \tag{33}
$$

Due to the normalizing step in pre-processing, we can assume the spread of per-sample means to be very small: $\frac{1}{n}\sum_{i=1}^n (s_i - \overline{s})^2 = \frac{1}{n}\|d\|_2^2 \leq \rho^2$ for $\rho \in [0, 1]$, then we have

$$
|B^{(\beta)}| \leq \rho\sqrt{n}\|Y^{(\beta)}\|_2 \tag{34}
$$

Plugging (34) into (32), we complete the proof.

□

#### 5.4 Bound $\mu^{(\beta)}$ for A Fixed $\beta \in \mathbb{R}^{k\times 1}$

Now if we assume $Y^{(\beta)}$ to be good as in Definition 1, which then implies $\frac{\|Y^{(\beta)}\|_1}{\|Y^{(\beta)}\|_2} = \Theta(\sqrt{n})$. Along with Lemma 4, we can complete the proof of Lemma 1.

**Proof of Lemma 1.** Dividing (31) by $P^{(\beta)}$ defined in (18), we get for high probability $> 1 - \frac{0.01}{(3/\epsilon)^k}$,

$$
\frac{|Q^{(\beta)}|}{P^{(\beta)}} \leq \frac{\|Y^{(\beta)}\|_2}{\|Y^{(\beta)}\|_1} \Big(\sqrt{2 \ln\Big(\frac{2(3/\epsilon)^k}{0.01}\Big)} + \rho\sqrt{n}\Big) = \frac{\sqrt{2 \ln\Big(\frac{2(3/\epsilon)^k}{0.01}\Big)} + \rho\sqrt{n}}{R^{(\beta)}} \tag{35}
$$

where $R^{(\beta)} = \frac{\|Y^{(\beta)}\|_1}{\|Y^{(\beta)}\|_2} = \Theta(\sqrt{n})$ as shown in (25) from Section 5.2 if we assume good $Y^{(\beta)}$ as indicated in Definition 1. Then with high probability $> 1 - \frac{0.01}{(3/\epsilon)^k}$, we have

$$r^{(\beta)} = \frac{|Q^{(\beta)}|}{P^{(\beta)}} \leq O\Big(\frac{\sqrt{2\ln\left(\frac{2(3/\epsilon)^k}{0.01}\right)}}{\sqrt{n}}\Big) + O(\rho) \tag{36}$$

Recall $\mu^{(\beta)}$ definition from (21), we know

$$\frac{1 - r^{(\beta)}}{1 + r^{(\beta)}} \leq \mu^{(\beta)} = \frac{P^{(\beta)} + Q^{(\beta)}}{P^{(\beta)} - Q^{(\beta)}} \leq \frac{1 + r^{(\beta)}}{1 - r^{(\beta)}} \tag{37}$$

After rearranging, we have

$$|\mu^{(\beta)} - 1| \leq \frac{2r^{(\beta)}}{1 - r^{(\beta)}} \tag{38}$$

and finally conclude the proof. □

#### 5.5 Extension to All $\beta \in \mathbb{R}^{k\times 1}$ with $\epsilon$-net and Union Bound

In order to bound $\mu$, we need Lemma 1 for all possible $\beta$. But there is infinitely many $\beta$. So a usual union bound would not apply. Instead, we consider the so-called $\epsilon$-net of $\mathbb{S}^{k-1}$.

**Definition 2** ([7, Definition 4.2.1. $\epsilon$-net with Euclidean distance ])**.** *Let $K \subseteq \mathbb{R}^k$ and let $\epsilon \in (0,1)$. A set $\mathcal{N} \subseteq K$ is an* $\epsilon$-net of $K$ (in $\ell_2$) *if for every $v \in K$ there exists $w \in \mathcal{N}$ such that $\|v - w\|_2 \leq \epsilon$. Equivalently, $K \subseteq \bigcup_{w\in\mathcal{N}} B_2(w,\epsilon)$, where $B_2(w,\epsilon) = \{x \in \mathbb{R}^k : \|x - w\|_2 \leq \epsilon\}$. We will mainly apply this to $K = \mathbb{S}^{k-1}$.*

It is a well-known fact that there exists a good sized $\epsilon$-net of $\mathbb{S}^{k-1}$.

**Fact 2** ([7, Corollary 4.2.13. $\epsilon$-net for the sphere])**.** *For any dimension $k$ and any $\epsilon \in (0,1)$, there exists an $\epsilon$-net $\mathcal{N} \subset \mathbb{S}^{k-1}$ such that*

$$|\mathcal{N}| \ \leq \ \left(\tfrac{3}{\epsilon}\right)^k \tag{39}$$

*and for every $v \in \mathbb{S}^{k-1}$ there exists $w \in \mathcal{N}$ with*

$$\|v - w\|_2 \ \leq \ \epsilon \tag{40}$$

The usefulness of Fact 2 is that it reduces statements about all $\beta \in \mathbb{S}^{k-1}$ to statements about only the finitely many $\beta \in \mathcal{N}$. Indeed, if a bound holds uniformly for all $\beta \in \mathcal{N}$, then for an arbitrary $\beta \in \mathbb{S}^{k-1}$ one may pick a nearby $\hat\beta \in \mathcal{N}$ with $\|\beta - \hat\beta\|_2 \leq \epsilon$ and transfer the bound from $\hat\beta$ to $\beta$ up to an additional error term proportional to $\epsilon$. Thus, combining a union bound over $\mathcal{N}$ yields a high-probability guarantee for all $\beta \in \mathbb{S}^{k-1}$.

**Lemma 5** (Bound $Q^{(\beta)}$ for All $\beta \in \mathbb{R}^{k\times 1}$)**.** *Fix $\epsilon \in (0,1)$ and let $\mathcal{N}$ be an $\epsilon$-net of the unit sphere $\mathbb{S}^{k-1}$ with $|\mathcal{N}| \leq (3/\epsilon)^k$. Then with probability at least* 0.99*, for all $\beta \in \mathbb{S}^{k-1}$,*

$$|Q^{(\beta)}| \ \leq \ \|H'\beta\|_2\Big(\sqrt{2\ln\left(\tfrac{2(3/\epsilon)^k}{0.01}\right)} + \rho\sqrt{n}\Big) \ + \ \sqrt{n}\,\|H'\|_2\,\epsilon \tag{41}$$

*Proof.* For each $\hat\beta \in \mathcal{N}$, define the failure event

$$\mathcal{F}_{\hat\beta} := \left\{ |Q^{(\hat\beta)}| > \|Y^{(\hat\beta)}\|_2\Big(\sqrt{2\ln\Big(\tfrac{2(3/\epsilon)^k}{0.01}\Big)} + \rho\sqrt{n}\Big) \right\} \tag{42}$$

i.e. the complement of the event guaranteed in (31) from Lemma 4. By Lemma 4, each $\hat\beta \in \mathcal{N}$ has

$$\Pr(\mathcal{F}_{\hat\beta}) \ \leq \ \frac{0.01}{(3/\epsilon)^k} \tag{43}$$

Applying the union bound over all $\hat{\beta} \in \mathcal{N}$ then yields

$$\Pr\Big( \bigcup_{\hat{\beta}\in\mathcal{N}} \mathcal{F}_{\hat{\beta}} \Big) \;\le\; |\mathcal{N}| \cdot \frac{0.01}{(3/\epsilon)^k} \;\le\; 0.01 \tag{44}$$

Thus we can say, $\forall\, \hat{\beta} \in \mathcal{N}$, with probability $> 0.99$,

$$\forall\, \hat{\beta} \in \mathcal{N}: \quad |Q^{(\hat{\beta})}| \;\le\; \|H'\hat{\beta}\|_2 \Big( \sqrt{2 \ln\Big(\frac{2(3/\epsilon)^k}{0.01}\Big)} + \rho\sqrt{n} \Big) \tag{45}$$

For the rest of the $\beta \in \mathbb{S}^{k-1} \backslash \mathcal{N}$, pick a nearest net point $\hat{\beta} \in \mathcal{N}$ with $\|\beta - \hat{\beta}\|_2 \le \epsilon$. Since $\|y\|_2 \le \sqrt{n}$, and by applying triangle inequality, we have

$$|Q^{(\beta)}| \;\le\; |Q^{(\hat{\beta})}| + |y^\top H'(\beta - \hat{\beta})| \;\le\; |Q^{(\hat{\beta})}| + \|y\|_2 \, \|H'\|_2 \, \|\beta - \hat{\beta}\|_2 \;\le\; |Q^{(\hat{\beta})}| + \sqrt{n}\, \|H'\|_2 \, \epsilon \tag{46}$$

Combine (45) with (46) to conclude the stated bound for all unit $\beta$.

□

**Proof of Lemma 2.** Rearranging (41) from Lemma 5 and recall $Y^{(\beta)} = H'\beta$, we have

$$|Q^{(\beta)}| \;\le\; \|Y^{(\beta)}\|_2 \Big( \sqrt{2\big(\ln 200 + k \ln(3/\epsilon)\big)} + \rho\sqrt{n} \Big) \;+\; \sqrt{n}\, \|H'\|_2 \, \epsilon \tag{47}$$

Divide both sides of (47) by $P^{(\beta)} = \|Y\|_1$, then we know, for any unit $\beta$,

$$r^{(\beta)} \;=\; \frac{|Q^{(\beta)}|}{\|Y^{(\beta)}\|_1} \;\le\; \frac{\|Y^{(\beta)}\|_2}{\|Y^{(\beta)}\|_1} \Big( \sqrt{2(\ln 200 + k\ln(3/\epsilon))} + \rho\sqrt{n} \Big) \tag{48}$$

$$+\; \frac{\sqrt{n}\, \|H'\|_2 \, \epsilon}{\|Y^{(\beta)}\|_1} \tag{49}$$

Since the good $Y^{(\beta)}$ works for any $\beta$, therefore $\frac{\|Y^{(\beta)}\|_2}{\|Y^{(\beta)}\|_1} = \Theta(\frac{1}{\sqrt{n}})$.
In addition to $\frac{1}{\|H'\beta\|_1} \le \frac{1}{\|H'\beta\|_2} \le \frac{1}{\sigma_{\min}(H')}$, then for probability $> 0.99$,

$$r^{(\beta)} \le \sqrt{\frac{2(\ln 200 + k\ln(3/\epsilon))}{n}} + \rho \tag{50}$$

$$+\; \sqrt{n}\, \frac{\|H'\|_2}{\sigma_{\min}(H')}\, \epsilon \tag{51}$$

Taking the supremum over all unit $\beta$ yields (6) which concludes the proof.

□

#### 5.6 Definition 1 Holds for All $\beta \in \mathbb{R}^{k\times 1}$

Recall in Section 5.2, we proposed an assumption: More than $n/c$ entries in $Y^{(\beta)}$ have values greater than $t \cdot \frac{\|Y^{(\beta)}\|_2}{\sqrt{n}}$. In Claim 1, we proved that if such assumption holds, then we can get a good $Y^{(\beta)}$ as defined in Definition 1. Now for this section, we aim to show that Definition 1 holds if rows of $H$ are independently sampled for any $\beta$.

##### 5.6.1 $\ell_2$ Leverage Score

Towards the proof we need the following definition, and its useful properties.

**Definition 3** ($\ell_2$ Leverage Score)**.** *Denote the ith row of matrix $H \in \mathbb{R}^{n\times d}$ as $h_i$ for $i \in [n]$. The ith row $\ell_2$ leverage score for $H$ can be written as*

$$\tau_i(H) = h_i^\top (H^\top H)^{-1} h_i \tag{52}$$

Then we will need the following three facts and one lemma:

**Fact 3.** *Consider the SVD decomposition of $H \in \mathbb{R}^{n\times d}$ as $H = U\Sigma V^\top \in \mathbb{R}^{n\times d}$. Write $u_1, \dots, u_n$ as the rows of $U$. Then*

$$\tau_i(H) = \|u_i\|_2^2$$

*Proof.* From (52) and use $e_i$ as the standard basis vector, we get the following transformation

$$\begin{aligned}
\tau_i(H) &= e_i^\top H(H^\top H)^{-1}H^\top e_i \\
&= e_i^\top (U\Sigma V^\top)(V\Sigma^2V^\top)^{-1}(V\Sigma U^\top)e_i \\
&= e_i^\top U\Sigma\Sigma^{-2}\Sigma U^\top e_i \quad \text{(since } (V^\top V) = I \text{ and } (V\Sigma^2V^\top)^{-1} = V\Sigma^{-2}V^\top) \\
&= e_i^\top UU^\top e_i \qquad\qquad \text{(simplify } \Sigma\Sigma^{-2}\Sigma = I) \\
&= \|U^\top e_i\|_2^2 \qquad\qquad \text{(since } e_i^\top UU^\top e_i = \|U^\top e_i\|_2^2) \\
&= \|u_i\|_2^2
\end{aligned} \tag{53}$$

□

**Fact 4** (PCA Preserves $\ell_2$ Leverage Score)**.**

$$\tau_i(H) = \tau_i(H') \tag{54}$$

We leave verifying Fact 4 to the reader. $H' = U\tilde{\Sigma}$ where $\tilde{\Sigma}$ is the first $k$ values of $\Sigma$ (as $\Sigma$ is a diagonal matrix). Since we are using the same $U$ for both $H$ and $H'$, due to Fact 3, both values should match.

Finally, we need the fact that an invertible linear transformation also does not change the leverage score.

**Fact 5** (Whitening Preserves $\ell_2$ Leverage Score)**.** *Whitening linear transformation is irrelevant to our $\ell_2$ leverage scores. More formally, denote whitening as a square invertible matrix $W \in \mathbb{R}^{d\times d}$, then on our data $H \in \mathbb{R}^{n\times d}$, we have*

$$\tau_i(HW) = \tau_i(H) \quad \textit{for all } i \in [n] \tag{55}$$

*where $\tau_i(\cdot)$ represents the ith row $\ell_2$ leverage score.*

*Proof.* Recall the $\ell_2$ leverage score is defined as

$$\tau_i(H) = h_i^\top (H^\top H)^{-1}h_i = e_i^\top H(H^\top H)^{-1}H^\top e_i \tag{56}$$

where $e_i$ represents the standard basis vector. By multiplying the whitening matrix $W \in \mathbb{R}^{d\times d}$ on the right of our data $H \in \mathbb{R}^{n\times d}$, we have the following transformation steps:

$$\begin{aligned}
\tau_i(HW) &= e_i^\top (HW)((HW)^\top(HW))^{-1}(HW)^\top e_i \\
&= e_i^\top (HW)(W^\top H^\top HW)^{-1}(HW)^\top e_i \\
&= e_i^\top (HW)W^{-1}(H^\top H)^{-1}(W^\top)^{-1}(HW)^\top e_i \\
&= e_i^\top HWW^{-1}(H^\top H)^{-1}(W^\top)^{-1}W^\top H^\top e_i \\
&= e_i^\top H(H^\top H)^{-1}H^\top e_i \\
&= \tau_i(H)
\end{aligned} \tag{57}$$

□

**Lemma 6.** *$\ell_2$ leverage score for our data $H$ is near uniform. More formally, given that $H$ is generated by i.i.d. rows and underwent whitening operation (see Fact 5), using the same notations as above, with probability at least $1 - \exp(-\Theta(\epsilon^2 n))$ where $\epsilon \in [0, 1]$, $\ell_2$ leverage score for all $i \in [n]$ satisfies*

$$\frac{O(d)}{(1+\epsilon)n} \le \tau_i(H) \le \frac{O(d)}{(1-\epsilon)n} \tag{58}$$

*where $\tau_i(\cdot)$ represents the ith row $\ell_2$ leverage score.*

**Proof Sketch.** Let $H \in \mathbb{R}^{n\times d}$ with i.i.d. rows $h_i$. Write $X_i = h_i^\top h_i$, then we have $H^\top H = \sum_{i=1}^n X_i$. The following claim allows us to assume $\mathbb{E}[X_i] = I_d$ without loss of generality.

Due to Matrix Chernoff [8], with probability at least $1 - \exp(-\Theta(\epsilon^2 n))$,

$$(1-\epsilon)n\, I_d \;\preceq\; H^\top H \;\preceq\; (1+\epsilon)n\, I_d \tag{59}$$

Inverting yields

$$\frac{1}{(1+\epsilon)n} I_d \;\preceq\; (H^\top H)^{-1} \;\preceq\; \frac{1}{(1-\epsilon)n} I_d \tag{60}$$

Therefore we get

$$\frac{\|h_i\|^2}{(1+\epsilon)n} \;\leq\; \tau_i(H) \;\leq\; \frac{\|h_i\|^2}{(1-\epsilon)n} \tag{61}$$

Due to whitening, for all $i \in [n]$, the row norms $\|h_i\|^2$ concentrate around $d$. Then it follows that all leverage scores satisfy $\tau_i(H) \approx \frac{d}{n}$ i.e. near-uniform across rows.

□

##### 5.6.2 Complete the Proof

Now with the necessary components, we are ready to prove Lemma 7 shown below.

**Lemma 7** (Satisfiable)**.** *$H'$ satisfies Definition 1.*

*Proof.* A direct corollary from Fact 4, Fact 5 and Lemma 6 is the near-uniformity of $\ell_2$ leverage scores on our data matrix $H' \in \mathbb{R}^{n\times k}$ which underwent PCA but without whitening. This implies that $\tau_i(H) = \|u_i\|_2^2 \approx k/n$ for all $i \in [n]$, where no single row dominates. Therefore we can say that with high probability over $H$, there exists a constant $z \geq 1$ such that

$$\sqrt{\tau_i(H)} = \|u_i\|_2 \;\leq\; z\sqrt{\tfrac{k}{n}} \quad \text{for all } i \in [n] \tag{62}$$

Now fix any unit vector $w \in \mathbb{R}^k$ and set $x = Uw \in \mathbb{R}^n$. Then $\|x\|_2 = 1$ and each coordinate satisfies

$$|x_i| = |\langle u_i, w\rangle| \;\leq\; \|u_i\|_2 \|w\|_2 \;\leq\; z\sqrt{\tfrac{k}{n}} \tag{63}$$

**Claim 2.** *Let $B := z\sqrt{k}$. For any threshold $t \in (0, B)$,*

$$\#\Big\{i \in [n] : |x_i| \;\geq\; \tfrac{t}{\sqrt{n}}\Big\} \;\;\geq\;\; n \cdot \frac{1-t^2}{B^2 - t^2} \tag{64}$$

*Proof.* Define $B = z\sqrt{k}$. Recall that $x := Uw \in \mathbb{R}^n$ with $\|x\|_2 = 1$ and $|x_i| \leq B/\sqrt{n}$ for all $i$. Fix a threshold $t \in (0, B)$ and define

$$S := \Big\{i \in [n] : |x_i| < \tfrac{t}{\sqrt{n}}\Big\}, \qquad m := |S| \tag{65}$$

Then its complement $S^c$ has size $n - m$. We can bound the squared norm of $x$ as

$$\|x\|_2^2 = \sum_{i=1}^n x_i^2 = \sum_{i\in S} x_i^2 + \sum_{i\in S^c} x_i^2 \tag{66}$$

For indices in $S$ we have $|x_i| < t/\sqrt{n}$, so $x_i^2 \leq t^2/n$. Thus

$$\sum_{i\in S} x_i^2 \;\leq\; \frac{mt^2}{n} \tag{67}$$

For indices in $S^c$ we only know $|x_i| \leq B/\sqrt{n}$, so

$$\sum_{i \in S^c} x_i^2 \;\leq\; \frac{(n-m)B^2}{n} \tag{68}$$

Combining (67) and (68) gives

$$1 = \|x\|_2^2 \;\leq\; \frac{mt^2}{n} + \frac{(n-m)B^2}{n} \tag{69}$$

Rearranging this inequality yields

$$m \;\leq\; n \cdot \frac{B^2 - 1}{B^2 - t^2} \tag{70}$$

Therefore the number of indices with $|x_i| \geq t/\sqrt{n}$ is

$$n - m \;\geq\; n - n \cdot \frac{B^2 - 1}{B^2 - t^2} = n \cdot \frac{1 - t^2}{B^2 - t^2} \tag{71}$$

Hence we conclude

$$\#\Big\{ i \in [n] : |x_i| \;\geq\; \tfrac{t}{\sqrt{n}} \Big\} \;\;\geq\;\; n \cdot \frac{1 - t^2}{B^2 - t^2} \tag{72}$$

□

Next, for any coefficient vector $\beta \in \mathbb{R}^k$, define $g = \Sigma V^\top \beta$ and $Y^{(\beta)} = Ug \in \mathbb{R}^n$. Write $w = g/\|g\|_2$ so that $x = Uw$ and $\|g\|_2 = \|Y^{(\beta)}\|_2$. Then each coordinate satisfies

$$Y_i \;=\; u_i^\top g \;=\; \|g\|_2 \,\langle u_i, w\rangle \;=\; \|Y\|_2 \, x_i \tag{73}$$

Therefore, according to Claim 2, we have

$$\#\Big\{ i \in [n] : |Y_i| \;\geq\; t \cdot \tfrac{\|Y\|_2}{\sqrt{n}} \Big\} \;\;\geq\;\; n \cdot \frac{1 - t^2}{B^2 - t^2} \tag{74}$$

This shows that for every choice of $\beta$ in the PCA subspace, at least a constant fraction of the coordinates of $Y$ lie above the threshold $t \cdot \|Y\|_2/\sqrt{n}$, which is exactly the assumption as seen in (26) from Definition 1.

□

### References


1. Dexter, G., Khanna, R., Raheel, J. & Drineas, P. *Feature Space Sketching for Logistic Regression* arXiv:2303.14284 [cs, stat]. Mar. 2023. (2024).
2. Eckart, C. & Young, G. The approximation of one matrix by another of lower rank. en. *Psychometrika* **1,** 211–218. ISSN: 0033-3123, 1860-0980. (2024) (Sept. 1936).
3. Cohen, M. B. & Peng, R. *$\ell_p$ Row Sampling by Lewis Weights* arXiv:1412.0588 [cs, math]. Dec. 2014. (2024).
4. Mai, T., Musco, C. N. & Rao, A. *Coresets for Classification – Simplified and Strengthened* en. in *Advances in Neural Information Processing Systems* (Nov. 2021). (2024).
5. Munteanu, A., Schwiegelshohn, C., Sohler, C. & Woodruff, D. *On Coresets for Logistic Regression* in *Advances in Neural Information Processing Systems* **31** (Curran Associates, Inc., 2018). (2024).
6. Hoeffding, W. Probability Inequalities for Sums of Bounded Random Variables. en. *Journal of the American Statistical Association* **58,** 13–30. ISSN: 0162-1459, 1537-274X. (2025) (Mar. 1963).
7. Vershynin, R. *High-Dimensional Probability: An Introduction with Applications in Data Science* 1st ed. ISBN: 978-1-108-23159-6 978-1-108-41519-4. (2024) (Cambridge University Press, Sept. 2018).
8. Tropp, J. A. User-friendly tail bounds for sums of random matrices. *Foundations of Computational Mathematics* **12.** arXiv:1004.4389 [math], 389–434. ISSN: 1615-3375, 1615-3383. (2025) (Aug. 2012).
